# The DNA methylation landscape of multiple myeloma shows extensive inter- and intrapatient heterogeneity that fuels transcriptomic variability

**DOI:** 10.1101/2020.10.01.321943

**Authors:** Jennifer Derrien, Catherine Guérin-Charbonnel, Victor Gaborit, Loïc Campion, Magali Devic, Elise Douillard, Nathalie Roi, Hervé Avet-Loiseau, Olivier Decaux, Thierry Facon, Jan-Philipp Mallm, Roland Eils, Nikhil C. Munshi, Philippe Moreau, Carl Herrmann, Florence Magrangeas, Stéphane Minvielle

## Abstract

**Background:** Cancer evolution depends on epigenetic and genetic diversity. Historically, in multiple myeloma (MM), subclonal diversity and tumor evolution have been investigated mostly from a genetic perspective.

**Results:** Here, we combined the notions of epipolymorphism and epiallele switching to analyze DNA methylation heterogeneity in MM patients. We show that MM is characterized by the continuous accumulation of stochastic methylation at the promoters of development-related genes. High entropy change is associated with poor outcomes and depends predominantly on partially methylated domains (PMDs). These PMDs, which represent the major source of inter- and intrapatient DNA methylation heterogeneity in MM, are linked to other key epigenetic aberrations, such as CpG island (CGI)/transcription start site (TSS) hypermethylation and H3K27me3 redistribution as well as 3D organization alterations. In addition, transcriptome analysis revealed that intratumor methylation heterogeneity was associated with low-level expression and high variability.

**Conclusion:** We propose that disordered methylation in MM is responsible for high epigenetic and transcriptomic instability allowing tumor cells to adapt to environmental changes by tapping into a pool of evolutionary trajectories.

## Background

Multiple myeloma (MM) is a neoplasm of plasma cells (PCs) with an incidence rate of approximately 5/100,000 in Europe. The median survival time of patients has improved substantially over the past decade. This is due to the establishment of high-dose therapy followed by autologous stem cell transplantation as a routine procedure, significant improvements in supportive care strategies, and the introduction and widespread use of drugs including immunomodulatory drugs, proteasome inhibitors, histone deacetylase inhibitors and monoclonal antibodies. Nevertheless, almost all patients ultimately relapse due to the emergence of more aggressive sub-populations of myeloma PCs resistant to therapeutic agents.

Several mechanisms have been suggested to explain the capacity of the subpopulations of myeloma cells within an individual to survive the pressure of frontline therapy and proliferate. These mechanisms include: the emergence of myeloma cells that achieve bortezomib resistance by decommitment from immunoglobulin synthesis [1] or by the derepression of growth factor receptors typically not associated with the plasma cell lineage [2], somatic mutations that emerge during disease progression involving key driver genes in MM such as the mono- or biallelic loss of TP53 [3, 4] or the biallelic loss of TRAF3 [5]. These mechanisms facilitate the expansion of proliferative subclonal populations. Genetic intratumor heterogeneity increases the evolutionary fitness potential of a rare subset of myeloma cells harboring a particular combination of molecular aberrations to survive when challenged by multiagent chemotherapy [4]. The disease evolves predominantly through a Darwinian process of clonal expansion, and the population of tumor PCs represents an admixture of competing genetic subclones [6, 5, 7, 8]. In most patients, the treatment pressure causes the profound reorganization and diversification of subclonal populations with complex dynamics of tumor evolution raising the possibility of biologically and clinically important cross-talk between subclones [9]. However, the small number of genetic alterations detected in MM and relevant in relapse mechanisms do not alone explain the profound phenotypic variability across patients [10, 4]. Beyond genetic diversity, other processes generate the intratumoral functional heterogeneity of cancer cells, including global epigenetic changes [11].

To address this question in MM, we conducted an analysis of the DNA methylation landscape of 47 myeloma samples from 26 patients. This study was motivated by the facts that (1) MM is characterized by extreme heterogeneity in median DNA methylation levels compared with normal plasma cells (NPCs) [12], (2) intratumor methylome variability is higher in various cancer types including other B-cell malignancies [13, 14, 15, 16], than in normal cells, and (3) genome-wide analysis of DNA methylation at single-base-pair resolution combined with analysis of the methylation states of neighboring CpGs (in the local sequence context) is now available [17].

## Results

### Intrapatient epigenetic heterogeneity arises from stochastic DNA methylation

To explore the global DNA methylation spectrum of MM patients, we compared whole-genome bisulfite sequencing (WGBS) datasets of myeloma cells from patients with those of controls (Additional file 1: Table S1). We found a marked global hypomethylation (Fig. 1a), as previously reported in MM and other cancers [12, 13, 15]. Individual CpG sites showed a predominantly bimodal pattern in MM patients (Fig. 1b) with methylation levels depending on the local density of CpG (Fig. 1c and Additional file 2: Table S2). Although MM displayed no evidence of aberrant DNA methylation at many genes (Additional file 3: Figure S1), we observed slight methylation gain at CpG islands (CGIs) and methylation loss at adjacent shores and shelves regions compared to NPCs (Fig. 1d), as illustrated in the genomic region of *DOC2B* (Fig. 1e).

**Figure 1.**
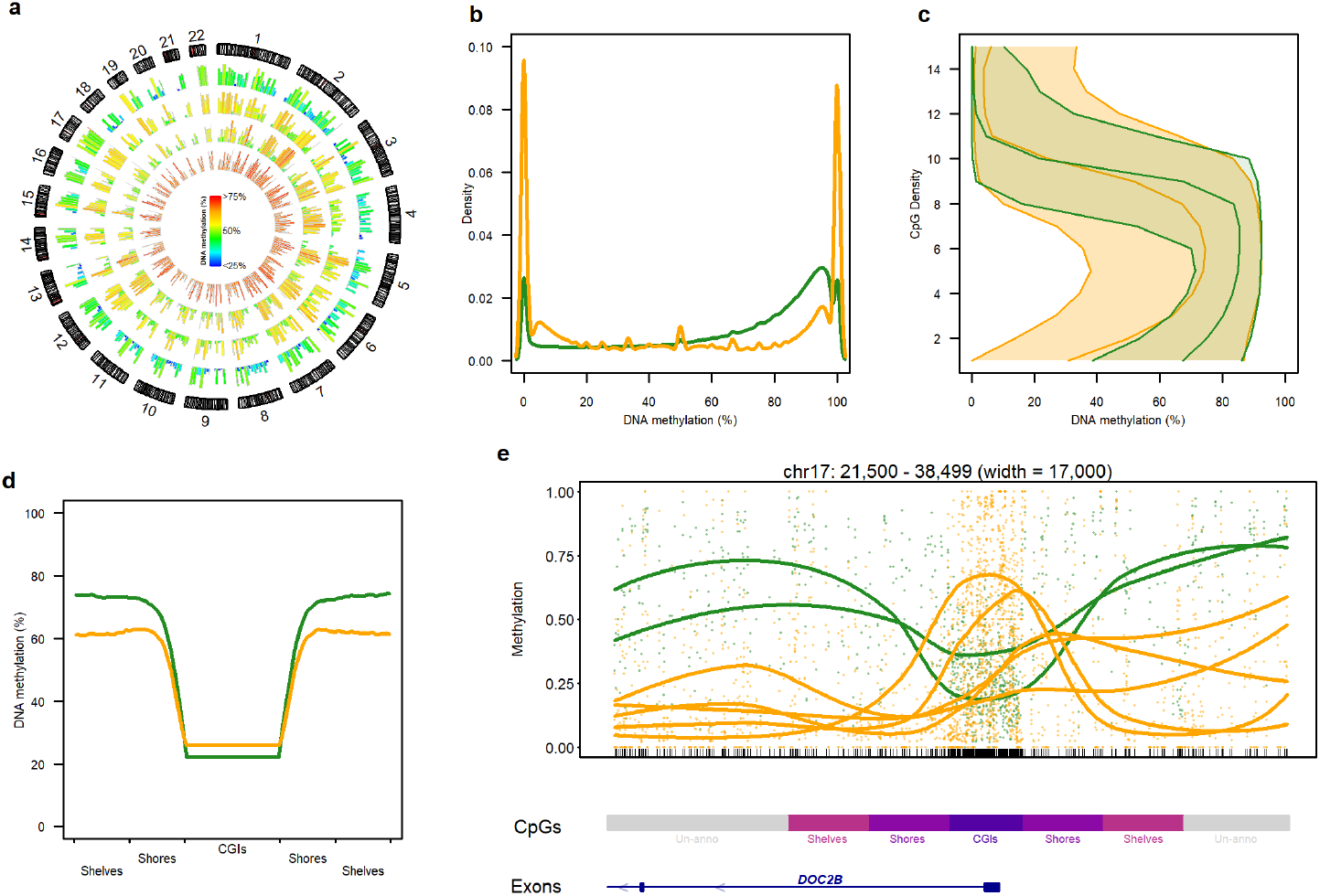
Global analysis of the DNA methylome of myeloma samples at diagnosis and NPC samples. (a) Circos plot of DNA methylation levels measured in WGBS data. From the center to the outside: NPC5 (data from the BLUEPRINT project, ERX301127), M#17, M#19, and M#10. Histograms represent CpG methylation levels averaged in 10-Mbp genomic windows. (b) Density distribution of DNA methylation levels measured in WGBS data for NPC (green, data from the BLUEPRINT project, ERX301127,ERX715130) and MM diagnosis (orange) samples. This color code was used in all figures. (c) CpG methylation within 100 bp genomic units as a function of local CpG density. (d) Mean methylation of CGIs and flanking regions in MM patients compared to NPCs. (e) Dispersive DNA methylation surrounding the *DOC2B* promoter in MM samples and in NPCs.

These results show that the MM epigenome is characterized by global DNA hypomethylation and variable intermediate methylation states depending on the local CpG content. We investigated whether these epigenetic changes could be a source of intrapatient heterogeneity. To assess the degree of epigenetic variability within patients, we employed enhanced reduced representation bisulfite sequencing (eRRBS) technology on 21 MM patients because eRRBS provides an average depth of 50X per covered CpG and approximately 2.7 million CpGs per sample, with more than 10X sequencing coverage (Additional file 4: Table S3 and Additional file 5: Table S4), and is cost effective for a large number of samples (48 in our study, Additional file 1: Table S1 and Additional file 3: Figure S2). We then applied a computational method that analyzes DNA methylation modifications at a genomic locus defined as a group of four contiguous CpGs covered by single sequence reads [17]. We computed for each genomic locus the epipolymorphism, defined as a probability distribution of the 16 possible methylation patterns (epialleles): a high value of epipolymorphism indicates a stochastic process of DNA methylation [18] (Additional file 3: Figure S3). Almost all genomic loci were located in CGIs (85% in MM samples) and, to a lesser extent, in promoter transcription start sites (TSSs) and gene bodies (74% in MM samples, Additional file 3: Figure S4). We plotted epipolymorphism distribution as a function of methylation level for all loci in NPCs and MM samples (Fig. 2a). Remarkably, loci with modest methylation levels (5% - 25%) showed a higher degree of epipolymorphism in cancer cells than in control cells whereas at higher methylation levels, epipolymorphism distribution was similar in MM and NPCs, indicating enrichment of a fraction of loci-CGIs with a stochastic methylation state at the expense of a bimodal methylation profile in MM samples. In all cases, promoter CGI methylation gain was associated with an increase in epipolymorphism (Additional file 3: Figure S5).

**Figure 2.**
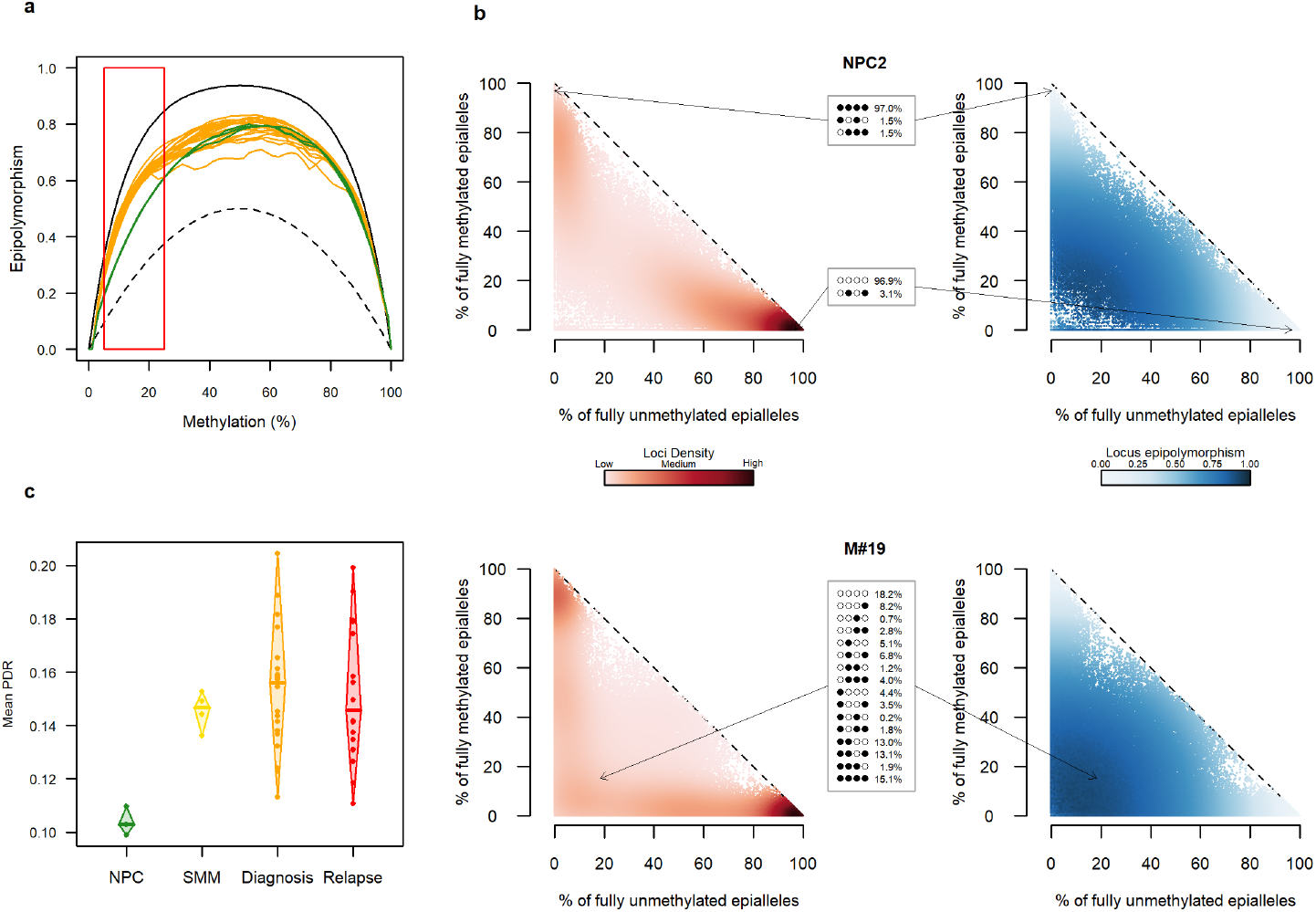
Intrapatient DNA methylation heterogeneity in MM. (a) Epipolymorphism levels as a function of the average DNA methylation at each locus in NPC (green) and diagnosis (orange) samples. The maximal epipolymorphism (continuous black line) and the bimodal epipolymorphism (dotted black line) are represented. (b) Scatterplots showing loci organization for NPC2 and M#19 diagnosis samples. Each point corresponds to a locus of 4 adjacent CpGs. On the left, each point is color coded according to the density of the surrounding points; on the right, each point is color coded according to its epipolymorphism level. (c) Mean proportion of discordant reads per sample.

To further characterize epiallele patterns of each sample, we stratified loci according to the frequency of fully methylated or fully unmethylated epialleles (Fig. 2b and Additional file 3: Figure S6). NPC loci displayed near-bimodal distribution, with a large number of homogeneous loci either fully unmethylated or fully methylated (Fig. 2b, top). Conversely, in MM samples, a large number of loci showing high epipolymorphism levels were along the axes, indicating marked enrichment for stochastic (i.e., neither fully methylated nor fully unmethylated) methylation patterns (Fig. 2b, bottom). Notably, this intrasample methylation heterogeneity was already present at the early stage of the disease, recognized as smoldering MM (SMM), and persisted at relapse (Additional file 3: Figure S6). A complementary approach that measures the proportion of discordant reads (PDR) [15] confirmed the increase in epigenetic heterogeneity in MM cells compared to NPCs. We found that the average PDR (i.e. genomic loci with a large number of reads that contain both unmethylated and methylated CpGs over the total number of reads) was significantly higher (p-value < 0.05 (DTK pairwise)) in MM samples, regardless of the disease stage, than in NPCs (Fig. 2c).

Overall, these results revealed that during disease initiation, growth and progression, malignant PCs accumulate randomness in DNA methylation at the expense of a more coherent methylation state, leading to a high degree of intratumoral epigenetic heterogeneity in MM patients.

### Entropy changes in MM development

Since we have shown that stochastic DNA methylation gains lead to intratumor heterogeneity, we sought to analyze epiallelic dynamics during MM development and progression. We used an algorithm that allows the identification of epigenetic loci (eloci) that have a significant epiallele composition change (entropy change) between two states (e.g., normal vs cancer or diagnosis vs relapse). To compare global methylome changes across patient samples, the number of eloci was normalized according to the number of covered loci, referred to as eloci per million loci covered (EPM) (Additional file 3: Figure S7)[17]. We first examined the link between the degree of epiallelic shifting and outcome. For this purpose, patients were separated into two groups according to their EPM values. We found that MM patients with high EPM had reduced relapse-free survival (p-value = 0.02 (log-rank test); Fig. 3a). EPM was independent of age, sex and ploidy. Remarkably, the extent of entropy changes varied greatly between patients at diagnosis and persisted at relapse compared to NPCs (Fig. 3b, c). The genome-wide distribution of eloci showed a predominant location at CGIs (average 54% in MM vs NPCs). As a control, we determined eloci by comparing NPCs to NPCs (NPC eloci) and found, on average, 19% of eloci in CGIs (p-value (Wilcoxon) = 0.0018) and at promoter-TSS (average 15% in MM vs NPCs and average 3% in NPC eloci; p-value (Wilcoxon) = 0.0123) (Additional file 3: Figure S8). We observed a similar distribution of eloci in relapse samples compared to NPCs (p-value (Wilcoxon) = 0.75 (CGIs) and 0.82 (promoter-TSS), Additional file 3: Figure S8).

**Figure 3.**
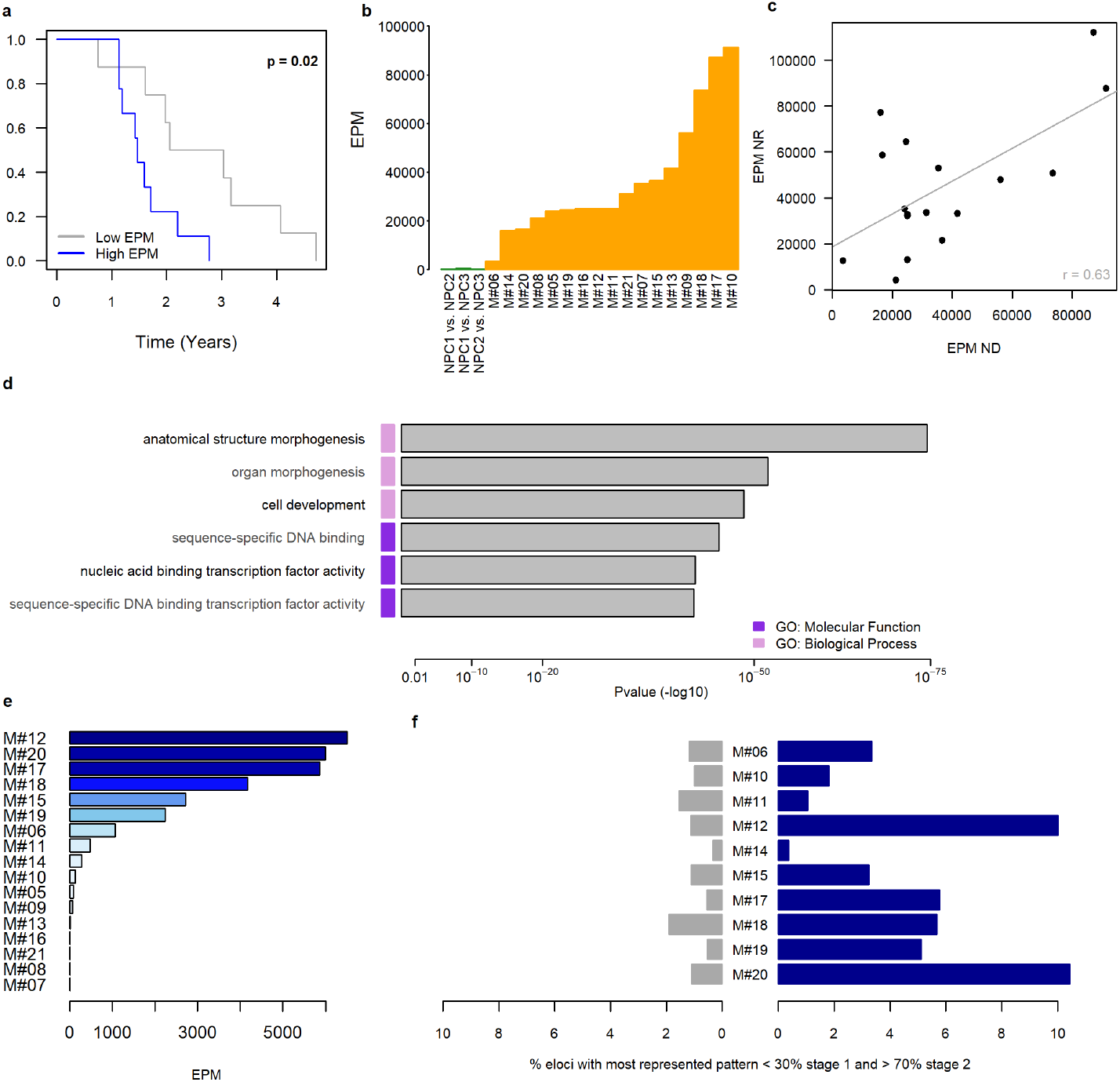
Evolution of epiallelic changes during the progression of MM and their clinical impact. (a) Time to relapse analysis for patients with high (blue, n=9) or low (gray, n=8) EPM values. (b) EPM values of NPC samples (NPC eloci, in green) and MM samples at diagnosis (MM eloci in orange). (c) Correlation between EPM at diagnosis and relapse compared to that in NPCs. (d) Enrichment analysis of eloci (NPC vs diagnosis) located in promoter CGIs and their associated ontological terms was performed with the GREAT tool [77]. (e) EPM value for each pairwise comparison between diagnosis and relapse (diagnosis vs relapse eloci). (f) Proportion of eloci presenting a selection of a methylation pattern at diagnosis compared to NPCs (in gray) and at relapse compared to diagnosis (in blue).

To gain insights into the biological functions of epigenetic perturbations, we performed gene ontology enrichment analysis for genes having eloci (MM diagnosis vs. NPCs) in promoter CGIs (Fig. 3d). Interestingly, associated genes showed marked enrichment for developmental regulators and transcription factor activity, while genes with promoter CGIs, including NPC eloci, were not enriched in any pathways or gene ontology terms. The identification of key signaling pathways rather than specific genes was not surprising considering the high DNA methylation variability across patients. Similar finding were reported by Easwaran et al. across different tumor types [19].

To further examine the impact of treatment on entropy changes, we assessed the epiallele shifting level during time to relapse by comparing diagnosis vs relapse paired samples (Fig. 3e). The degree of difference was highly variable from one patient to another; 24% of patients showed no epigenetic changes (EPM = 0) while 42% showed substantial changes (EPM > 1000) between diagnosis and relapse. We then focused our analysis on the evolution of extreme epiallele patterns (i.e., one predominant epiallele or a mixture of low represented epialleles) during two disease stages (i.e., before and after treatment). For this purpose, we compared epiallele shifts between NPCs and diagnosis samples and between paired samples at diagnosis and relapse. Interestingly, among the four possible epiallele pattern changes (Additional file 3: Figure S9), the selection pattern was significantly enriched at relapse compared to that at diagnosis (p-value (paired Wilcoxon) = 0.0059, Fig. 3f). Although this pattern involved a small number of eloci (median value = 4.2%), this finding suggests that treatment escape is associated with clonal selection at specific genomic loci.

Given the central role of chromatin in stabilizing gene expression and cellular states, we wanted to determined whether chromatin states are associated with genomic loci showing significant entropy changes in MM. To answer this question, we analyzed the distribution of eloci using 127 cell/tissue types at 25-state chromatin state segmentation [20] (Additional file 3: Figure S10a). Hypomethylated eloci were found in quiescent states while hypermethylated epiallele shifts were predominant in bivalent promoter-associated chromatin states (Additional file 3: Figure S10b). In addition, the frequency of these two subtypes of eloci varied widely between patients (Additional file 3: Figure S11).

### Abnormal bivalent promoter eloci CGI methylation increases intra- and intertumor heterogeneity and targets Wnt and Ras/MAPK signaling

We found that eloci were predominantly enriched (82.5%) in the bivalent promoters of embryonic stem cells (ESCs) (Additional file 3: Figures S10a and S12). In the embryonic system, bivalent promoter genes are not regulated by DNA methylation but rather by the simultaneous presence of the repressive mark H3K27me3 and the active transcription mark H3K4me3, which allow low basal transcription states that are dynamically inducible to ensure a balance between self-renewal and lineage commitment [21]. We compared the average change in the DNA methylation of eloci between MM samples and NPCs. In patient M#17, who had the highest number of eloci, we observed that 93% of the eloci acquired extensive gains of DNA methylation (higher than 40%) during neoplastic transformation (Fig. 4a and Additional file 3: Figure S13). Methylation gains are maintained during progression (Fig. 4a). This perturbation was associated with a decrease in gene expression in diagnosis and relapse samples compared to NPCs, whereas upregulated genes were present mostly at the time of diagnosis in almost all patients (Fig. 4b and Additional file 3: Figure S14). To further examine the relationship between gene expression and entropy changes in MM patients, we compared the expression levels between bivalent genes containing hypermethylated eloci in their promoter and bivalent genes without eloci. We found that disrupted (i.e., containing hypermethylated eloci) bivalent genes were less expressed but displayed greater interpatient variability (Fig. 4c; only genes with a mean expression level above 1 were taken into account for the coefficient of variation). Gene set enrichment analysis revealed that disrupted bivalent genes were enriched for important cancer-related pathways, including the Wnt and MAPK signaling pathways (Fig. 4d).

**Figure 4.**
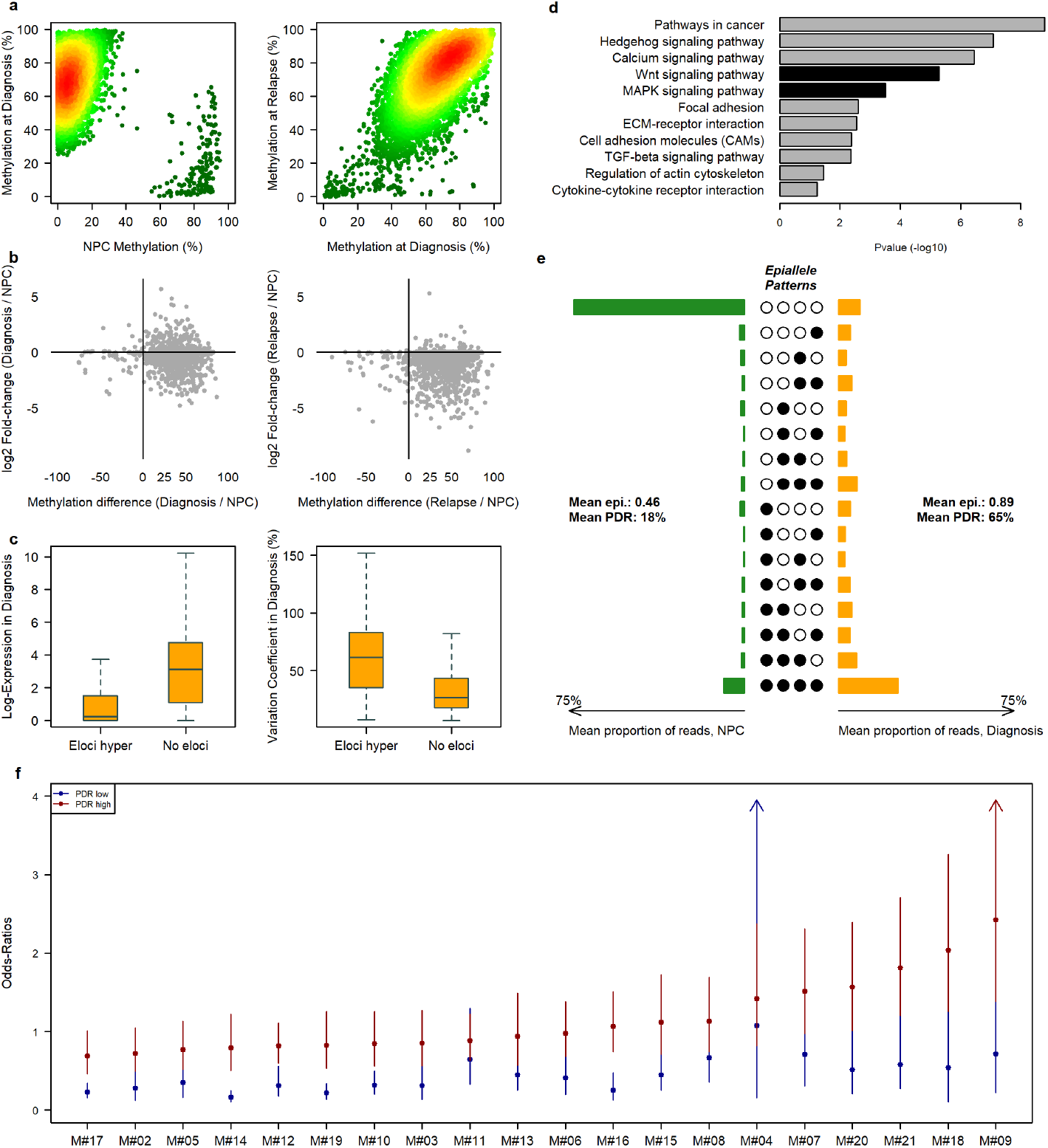
Methylation disruption in bivalent promoters in MM. (a) Scatterplot of eloci in bivalent promoters as a function of DNA methylation in NPCs and diagnosis (left) and as a function of DNA methylation at diagnosis and relapse (right) for patient M#17. The color gradient corresponds to the point density (low is green; high is red). (b) Scatterplot of DNA methylation of the eloci promoters versus RNA expression of the associated genes. (c) Normalized expression values of genes with a promoter CGI containing a hypermethylated elocus or no elocus (left) and the variation coefficient of these genes in MM samples (right). (d) Ontological analysis of genes with a bivalent promoter CGI affected by hypermethylated eloci. (e) Average methylation epiallele patterns of eloci in promoter CGIs for NPC and diagnosis samples (epi= epipolymorphism). (f) Odds ratios with 95% confidence intervals for the association between gene expression (FPM > 1) and promoter methylation (average methylation > 0.75/average methylation < 0.25) for genes with high (red) or low (blue) PDR levels in the promoter.

Myeloma cells are characterized by predominant stochastic methylation gains at bivalent promoters reflected by a high PDR and epipolymorphism, leading to intra-tumor heterogeneity (Fig. 4e). We then analyzed the consequences of intratumoral methylation variability on bivalent gene expression levels across the 20 MM samples. We separated the genes into two groups according to the PDR level of their bivalent promoter (lower or higher than the mean PDR) and calculated, in each group and for each patient, the odds ratio (OR) of the association between gene expression (FPM > 1 vs. *≤* 1) and bivalent promoter methylation (mean methylation < 25% vs. mean methylation > 75%; Fig. 4f and Additional file 6: Table S5). In all MM samples, promoters with a low PDR tended to maintain the expected opposite relationship between promoter methylation and transcription, whereas in promoters with a high PDR, for the majority of patients, the link between methylation and expression did not remain significant, and for some patients, we observed a significant link, but the relationship was opposite to the expected result (OR > 1). For example, *LIPG*, which showed comparable methylation levels in two samples (0.61 in M#14 and 0.63 in M#08) coupled with opposite expression levels (FPM of 4.57 in M#14 and 0.29 in M#08), demonstrates the decoupling relationship between promoter methylation and gene expression. M#14 displayed a high promoter PDR (0.80), whereas M#08 displayed a low promoter PDR (0.29, Additional file 3: Figure S15). OR analysis clearly showed the contribution of intratumoral methylation heterogeneity to increased transcriptional variability in MM. These results are concordant with those obtained in chronic lymphocytic leukemia (CLL) [15].

### Entropy changes towards hypomethylation preferentially occur in PMDs

Hypomethylated epiallele shifts are localized mainly in quiescent genomic regions (Additional file 3: Figure S10). We identified in most of the MM samples two subsets of hypomethylated eloci compared to normal samples (Fig. 5a, b and Additional file 3: Figure S16). The first type included extensively demethylated loci (Fig. 5b, bottom) and the second type contained epipolymorphic loci emerging due to partial methylation loss (Fig. 5b, top). Genomic regions that have lost their methylated state, termed partially methylated domains (PMDs), were initially discovered in a fibroblast cell line [22]. Several studies have reported cancer and noncancer human primary cells with PMDs [23, 24, 25, 26, 27]. PMDs cover approximately 50 to 75% of the genome of the human primary cell types and tissues investigated, while roughly a quarter are shared which indicates that PMDs retain strong tissue and cell type specificity characteristics [26]. To perform a comprehensive analysis of PMDs in MM, we used MethylSeekR [28] with default parameters to analyze WGBS data from our cohort of MM samples and the available WGBS dataset. We detected PMDs in both NPC samples and in only five MM samples from our cohort (Additional file 3: Figure S17). The PMD structure was highly similar between normal and MM cells despite a very variable level of DNA methylation (Additional file 3: Figure S18a). The base overlap was greater than 80%, the median length distribution was approximately 51 kb and the mean genome coverage was approximately 65% of the genome (Additional file 3: Figure S18b). Given the very good overlap between NPC and MM PMDs, we subsequently studied PMDs determined from NPCs (referred to as PC-PMDs) in MM patients.

**Figure 5.**
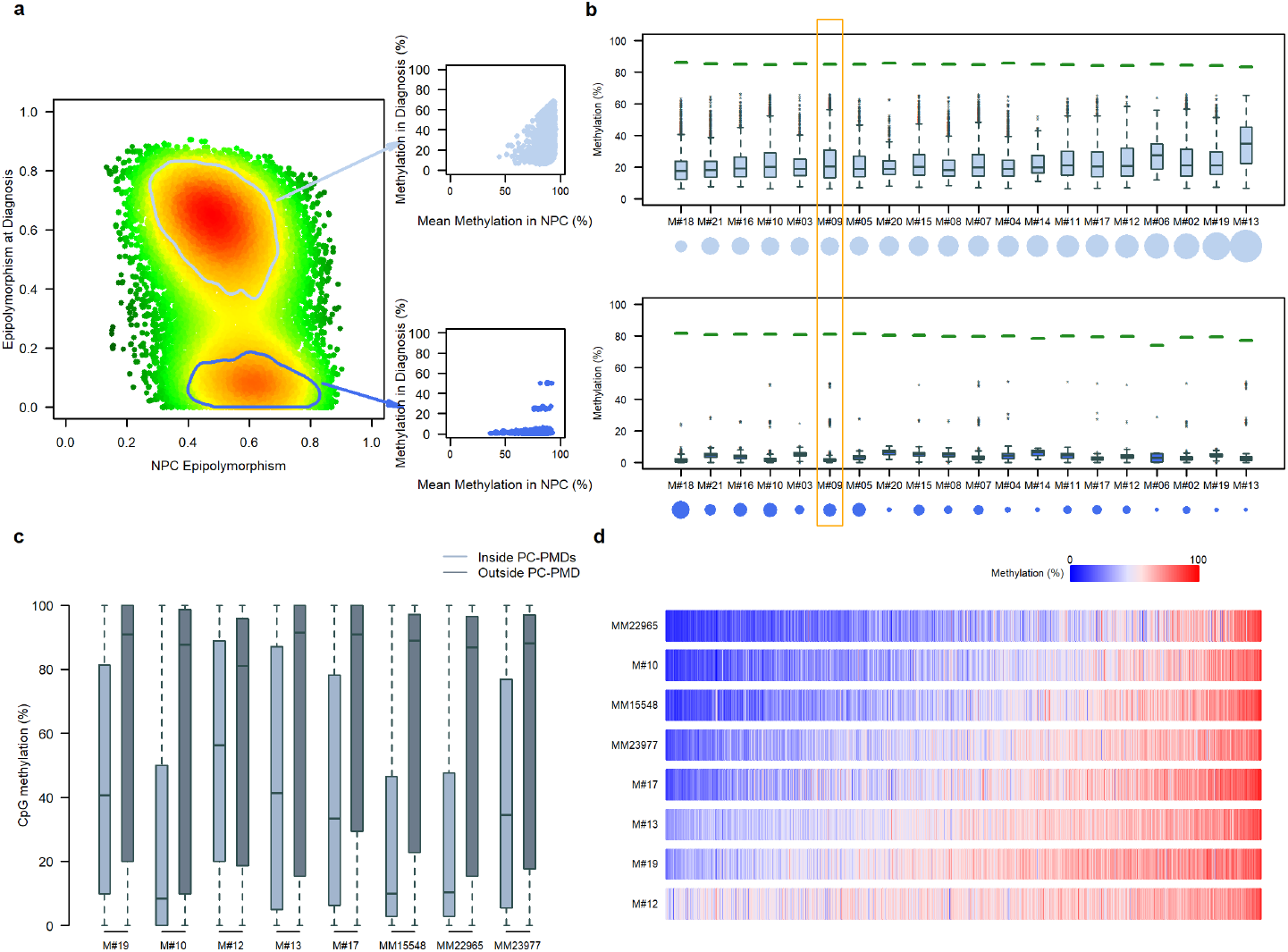
DNA methylation loss in partially methylated domains. (a) Scatterplot of hypomethylated eloci as a function of their epipolymorphism in NPC and diagnosis samples (M#09). The color gradient corresponds to the point density (low is green; high is red). Two eloci populations can be distinguished. The first population (top box) had a moderate decrease in methylation associated with a strong epipolymorphism, and the second population (bottom box) had a low heterogeneity and a drastic methylation decrease at diagnosis. (b) DNA methylation levels by patient of eloci with partial demethylation (eloci with an epipolymorphism value between 0.4 and 0.8 in NPCs and remain at the same level in MM) per patient are shown at the top, and DNA methylation levels of eloci with a decreased epipolymorphism value at diagnosis (eloci with an epipolymorphism value between 0.4 and 0.8 in NPCs, and an epipolymorphism value ≤0.2 at diagnosis) per patient are shown at the bottom. Green segments show the average methylation level of NPCs for these loci. The circles under the patient labels represent the size of the population compared to all hypomethylated eloci. Patient M#19, as an example in Fig. 5a, is outlined in orange. (c) DNA methylation levels of WGBS samples inside PC-PMD regions and outside PC-PMD regions (data from the BLUEPRINT project, see Additional file 1: Table S1). (d) Representation of the global DNA methylation of PC-PMDs in WGBS samples.

PC-PMD borders are not random and coincide with spatial genome organization, as indicated by the closeness of topological associated domain (TAD) borders to PC-PMD borders, which is larger than expected by chance (p < 0.001). In addition, PC-PMDs share key features with PMDs of various tumor types and normal tissues [24, 26, 27], including the correlation with lamina-associated domains (LADs) [29], late-replication timing and low gene density (Additional file 3: Figure S19a,b,c). Except for one sample, PC-PMDs that intersected with LADs displayed a long replication time and significantly low methylation levels (Additional file 3: Figure S19d,e). Importantly, the vast majority of hypomethylated eloci (70%) were located within PC-PMDs.

### Severe DNA methylation loss in PC-PMDs contributes to the redistribution of repressive histone marks and perturbations in CGIs/TSSs

We next examined the PC-PMDs methylation status across patients and the associated functional features. As expected, the DNA methylation level was lower within than outside of PC-PMDs (Fig. 5c) and variable between patients (Fig. 5d). In addition, this variability was associated with the PC-PMD length and replication time (Additional file 3: Figure S20). To study the relationship between DNA methylation and other epigenetic features, we integrated available WGBS data together with ChIP-seq data for histone marks. We found that the DNA methylation changes within PC-PMDs were associated with perturbations in both H3K9me3 and H3K27me3. Notably, the H3K9me3 deposit was associated with severe DNA methylation loss in long PC-PMDs (> 1 Mb) (Fig. 6a), as exemplified for the patient MM15548, with an unmethylated 3 Mb PC-PMD that was highly enriched with H3K9me3 (Fig. 6b). To our knowledge, such widespread H3K9me3 deposits associated with DNA methylation loss have so far been observed only in PMDs of cancer cell lines [26, 30, 31]. H3K27me3 was also perturbed inside PMDs. Notably, H3K27me3 enrichment was correlated with DNA methylation erosion (Fig. 6c). One notable example is the *DOCK3*-containing PC-PMD, which is enriched in H3K27me3 and showed H3K27me3 deposits in patients MM15548 and MM22965, who also exhibited DNA methylation erosion in this PMD; howerver, in another patient (MM23977) with a methylation level in this PMD comparable to that in NPCs, the H3K27me3 mark was absent (Fig. 6d). Taken together, these result show that perturbations in repressive histone marks are variable and depend on genomic regions, patients and DNA methylation levels.

**Figure 6.**
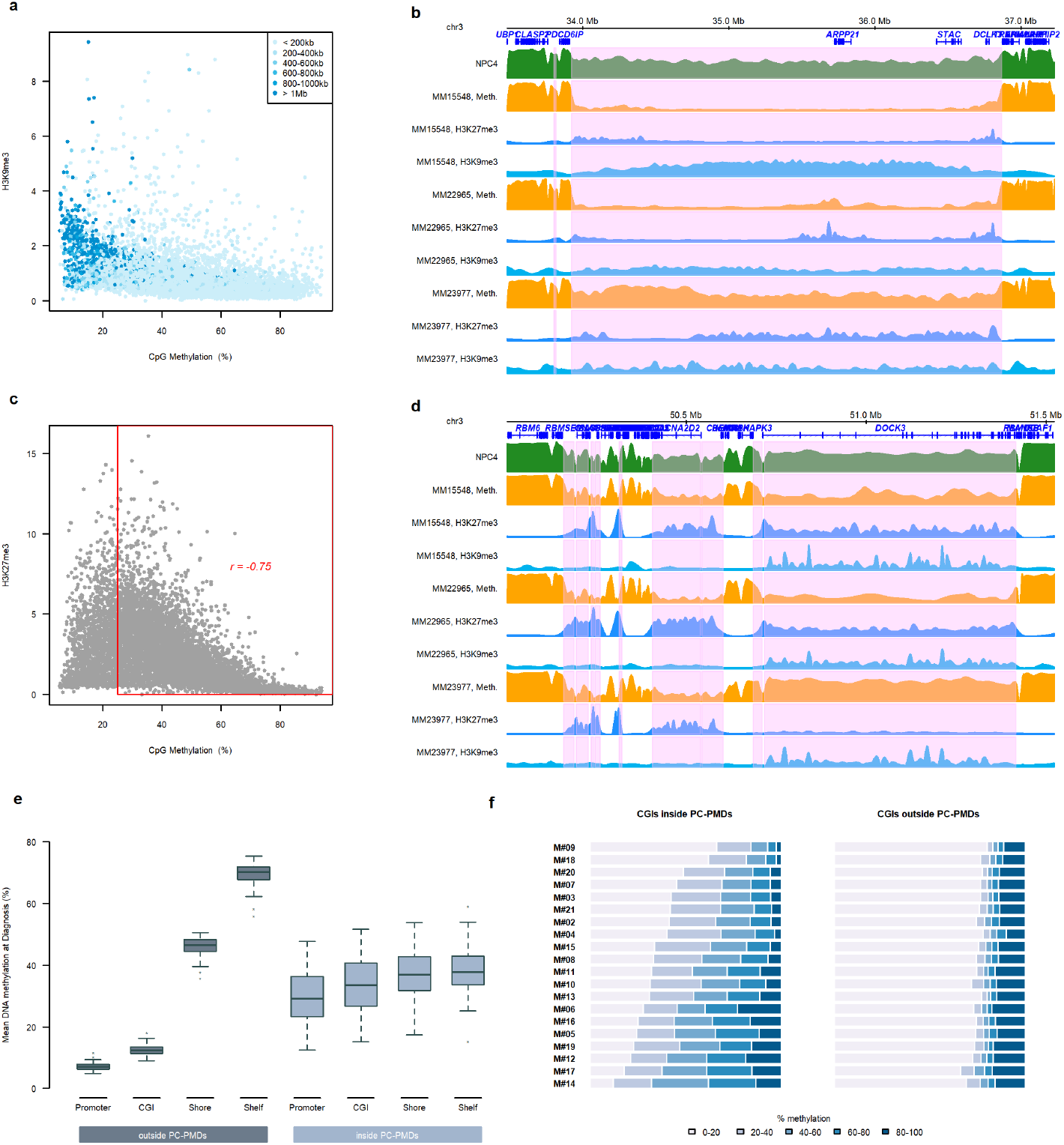
DNA methylation loss and associated DNA functional features. (a) The average methylation levels inside PC-PMDs according to average H3K9me3 intensity (data from the BLUEPRINT project: ERX1199099 and ERX712768). Each point represents a PC-PMD, and points are colored according to the size of the PC-PMD. (b) Example of a long unmethylated PC-PMD (pink area) associated with an increase in H3K9me3 in patient MM15548. (c) Average methylation levels in PC-PMDs according to their average H3K27me3 intensity (data from the BLUEPRINT project: ERX1199099 and ERX712769). (d) Example of PC-PMDs (pink areas) with methylation loss at diagnosis associated with an increase in H3K27me3 marks. (e) Mean DNA methylation level at diagnosis in promoter, CGIs, shores and shelves regions according to the presence or absence of PC-PMDs. (f) Proportion of CGIs with partial methylation inside and outside PC-PMDs.

We next investigated the impact of PC-PMD methylation erosion on regulatory regions. We found that the normal near-bimodal methylation state observed outside of PC-PMDs, was completely abolished inside PC-PMDs (Fig. 6e). As a result, promoter CGIs lost their hypomethylated state and gained intermediate methylation levels, while adjacent regions became less methylated to reach an intermediate degree of methylation. Notably, the acquisition of DNA methylation inside PC-PMDs resulted in a significant increase in the intermediate methylation state (>20% to <80%) of CGIs at the expense of strictly methylated or unmethylated states in all MM patients examined (Fig. 6f), in agreement with results obtained in breast cancer [24]. We noted that this phenomenon was also observed, albeit to a lesser extent, in NPCs (Additional file 3: Figure S21a); however, the percentage of loci with a high level of discordant reads (> 50%) was higher in all MM compared to normal samples (Additional file 3: Figure S21b) indicating that although PC-PMD methylation is perturbated in NPCs, CGI methylation patterns are more homogeneous (i.e., either fully methylated or unmethylated) (Additional file 3: Figure S21c).

Interestingly, a large proportion of bivalent promoter CGIs with disrupted DNA methylation was embedded within PC-PMDs (39% totally included and 54% with at least 30% of shared bases, as exemplified in Additional file 3: Figure S22). Furthermore, PC-PMD methylation loss was correlated with an overall epiallele shift increase (Spearman p = 0.46) (Additional file 3: Figure S23). These results support the notion that PC-PMD methylation loss may locally fuel epiallele shifts.

Given the interpatient heterogeneity of PMD methylation, we also investigated variations in gene expression. We found more gene expression variability inside PC-PMDs than outside of PC-PMDs; this difference was also maintained at relapse (Fig. 7a). Interestingly, we found that genes inside PC-PMDs were abundant within 186 kb of PC-PMD boundaries (Fig. 7b). We therefore focused our analysis on genes located near boundaries: Kyoto Encyclopedia of Genes and Genomes (KEGG) pathway analysis revealed specific enrichment in immune-related pathways including cytokine-cytokine receptor interaction, complement and coagulation cascades, autoimmune thyroid disease, natural killer cell-mediated cytotoxicity, antigen processing and presentation, regulation of autophagy, cell adhesion molecule and graft-versus-host disease (Fig. 7c). This enrichment pattern in PC-PMDs could provide an explanation for immune evasion mechanisms that occur during tumor progression [32, 33].

**Figure 7.**
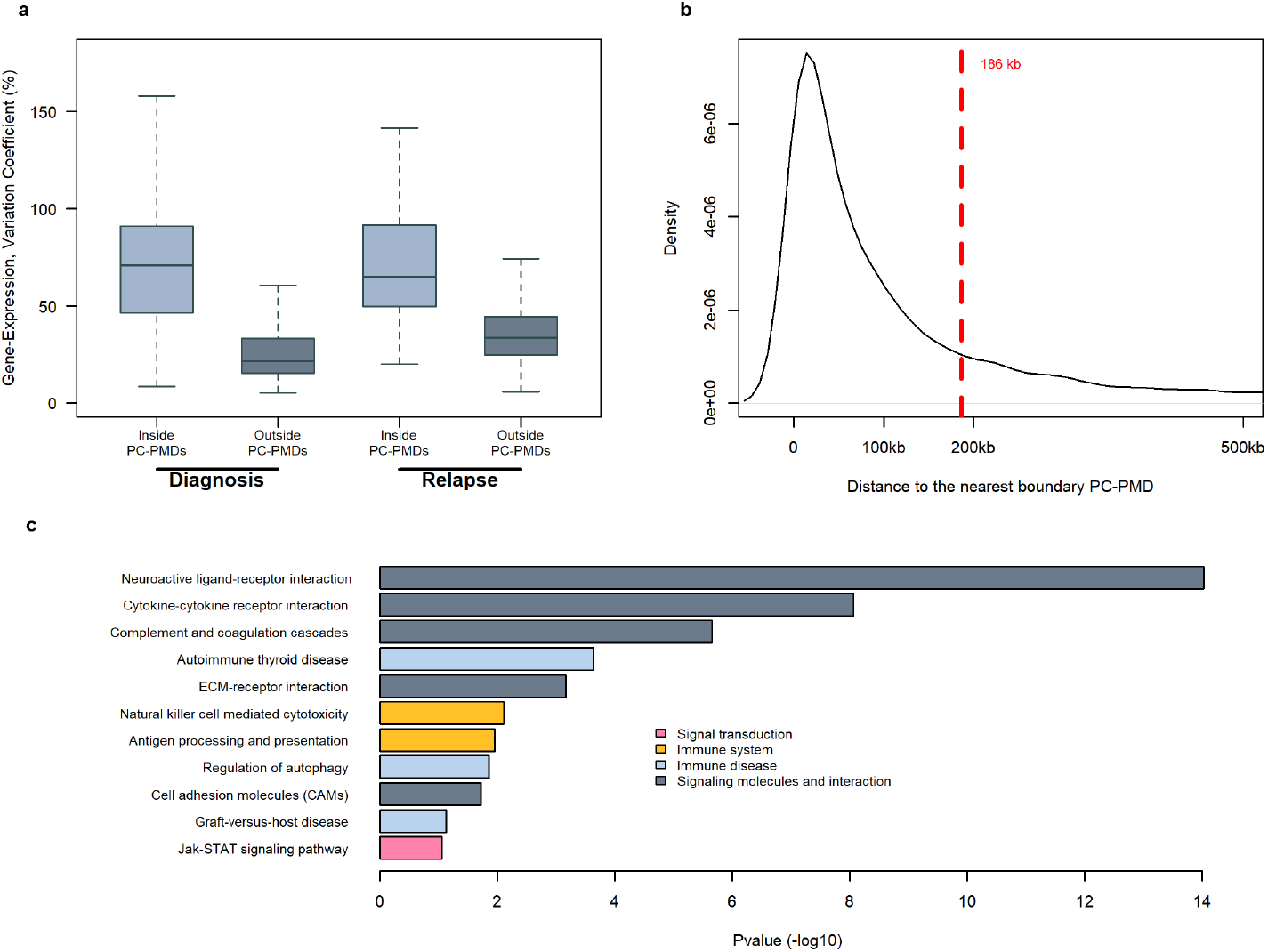
Gene expression variability in hypomethylated PC-PMDs. (a) Variation coefficient of gene expression in PC-PMD and non-PMD regions at diagnosis (left) and relapse (right). (b) Density of the TSS distance to the nearest PC-PMD boundary. The red dotted line corresponds to the 3rd quartile of the TSS distance to the nearest PC-PMD boundary. (c) Ontological analysis of genes in PC-PMDs located less than 186 kb from a PC-PMD boundary.

Taken together, these results show that MM displays a high epigenomic instability and great transcriptomic variability in PC-PMDs which might be beneficial for tumor cells.

### Three-dimensional (3D) reorganization is favored in hypomethylated PMDs

Global DNA methylation loss impacts spatial genome organization in the nucleus [34]. We wondered whether severe DNA hypomethylation that occurs during MM development could reshape 3D chromatin architecture in MM cells. The chromatin fiber of eukaryotic genomes is folded at multiple levels, including large-scale genomic structures to form distinct chromatin compartments A and B, characterized by gene-dense transcriptionally active open chromatin and gene-sparse transcriptionally closed chromatin, respectively [35]. During cancer development the genome is reorganized, and as a consequence, genomic regions of compartment A switch to those of compartment B and vice versa. The proportion of compartment A/B switching varies according to tumor type [36, 37]. We investigated the relationships between PC-PMDs and compartment A/B switching. We determined compartment A/B boundaries at 20-kb resolution from Hi-C data in a lymphoblastoid B cell line (GM12878), with a DNA methylome similar to that of PCs [12], and in the myeloma cell lines U266 and RPMI8226. We found that PC-PMDs were more prone to switch than other genomic regions (Fig. 8a). As a representative example, *IGF1R*, which encodes a major mediator of growth and survival in MM, and is located astride compartments A and B in NPCs, switched entirely to compartment A in both myeloma cell lines (Fig. 8b).

**Figure 8.**
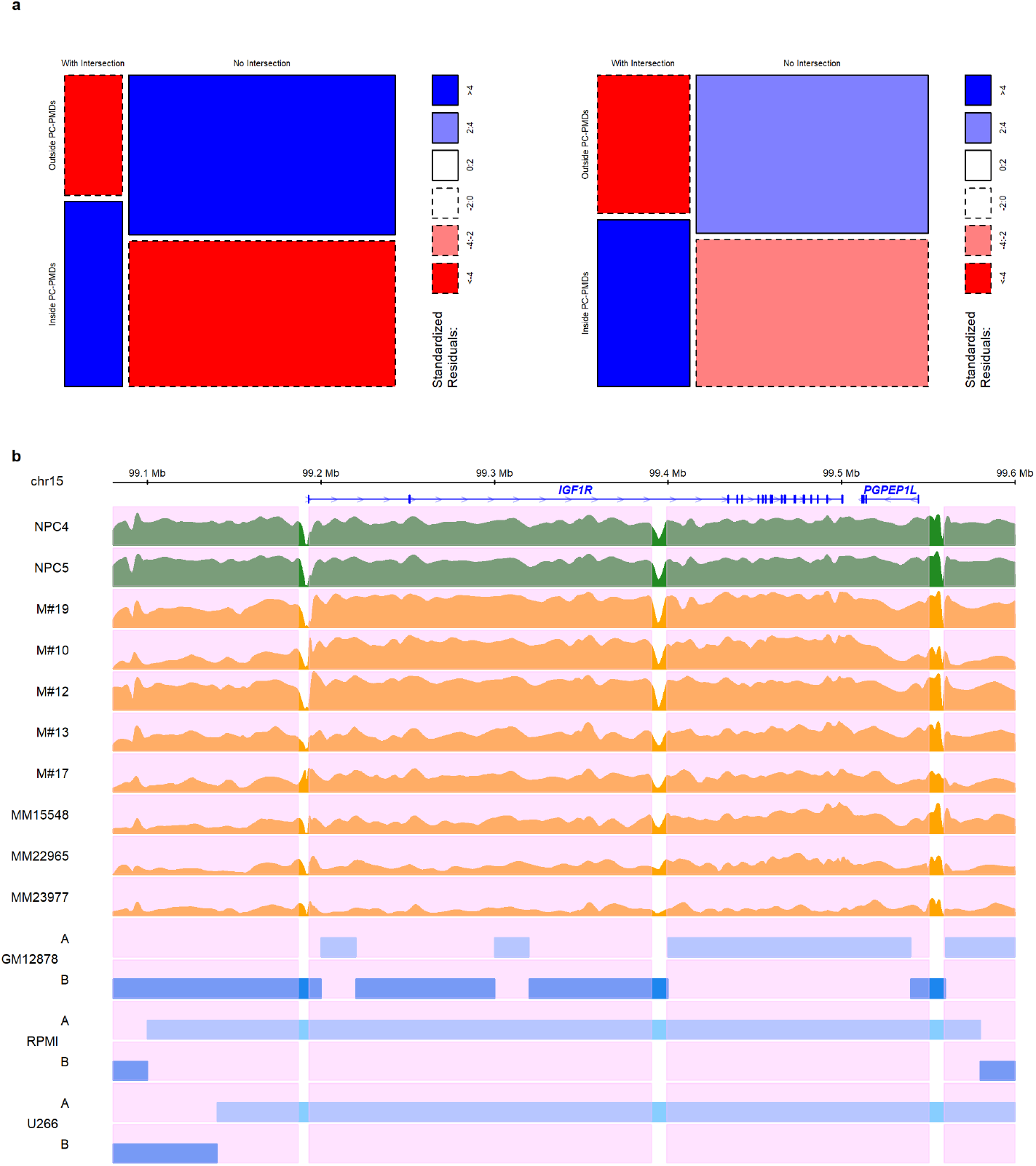
Hypomethylated PC-PMDs and the reorganization of compartment A/B. (a) Mosaïc plot illustrating the proportions of regions inside or outside PC-PMDs and with or without an intersection with a switch from B to A (left) and from A to B (right). (b) Example of *IGF1R* gene locus switching from compartment B to A (GM12878 cell line versus MM cell lines). Pink indicates PC-PMD regions.

Altogether, these results show that compartment switching might be favored by PMD instability.

## Discussion

MM is a heterogeneous disease. However, until now, only studies on genetic abnormalities have demonstrated the presence of subclonal populations that encountered complex evolution during the course of a disease. A recent study highlighted the contribution of intratumor epigenetic heterogeneity to shape CLL evolution after therapy [38]. To assess intrapatient DNA methylation heterogeneity in MM, we used the eRRBS technique, which allows us to capture the methylation status of individual CpGs in a single cell (more precisely in one single allele), with a large sequencing depth. Combining the notions of epipolymorphism and epiallele switching, we showed that the MM epigenome is characterized by high intratumor heterogeneity with a predominantly stochastic methylation pattern. Furthermore, a high methylation entropy change between malignant and NPCs is associated with a high risk of relapse.

We found that genes that undergo epiallele shifting are enriched in the control of developmental genes. Interestingly, in a model wherein hematopoietic progenitors are proposed to be the cells of origin in MM, an aberrant epigenetic program persisting through normal cell differentiation is implicated in tumor initiation [39]. Further analysis of this aberrant epigenetic program revealed strong enrichment of biological functions associated with developmental regulation (Additional file 3: Figure S24), suggesting that a disruption in developmental pathways could play a key role in the initiation of MM and increase susceptibility to oncogene transformation in response to environmental changes [40].

Entropy changes in bivalent promoters are associated with DNA methylation gains. These gains are stochastic, with an elevated PDR, which leads to a decoupling relationship between promoter methylation and transcription. These results are in line with those obtained in CLL [15]. Moreover, in CLL, single cell RNA-seq analysis showed that a high PDR is correlated with a “noisy” transcriptional landscape and an intermediate transcriptional state that interferes with complete silencing or high-level expression. Several studies have also shown the role of cellular heterogeneity and gene expression noise in overcoming drug resistance or metastatic barriers [41, 42, 43]. Together, these data suggest that increased stochastic methylation variation allows tumor cells to better adapt and find new trajectories in response to environmental changes.

A large majority of entropy changes between normal and malignant plasma cells associated with DNA methylation loss are found in regions that are already partially methylated in normal cells, indicating that PC-PMDs delimit genomic regions in which methylation information is not accurately maintained due to reduced energy consumption and channel capacity, as recently demonstrated for compartment B [44]. These large regions that cover approximately 70% of the genome are the major source of variability within MM patients. The PMDs of naive B cells display strong, progressive DNA methylation loss with differentiation into PCs [25]. During their maturation, PCs undergo successive rounds of division coupled with DNA methylation loss, thus supporting the idea that PMD hypomethylation is linked to mitotic dynamics [45, 46, 47].

In myeloma cells, PC-PMD hypomethylation is associated with other key epigenetic aberrations, such as promoter CGI hypermethylation. Indeed, we observed a loss of the hypomethylated state and gained intermediate methylation levels in regulatory regions within PC-PMDs. This disturbance has already been observed in NPCs; however, disordered methylation at promoters is lower in NPCs than in malignant cells, indicating a homogeneous methylation pattern at the promoter being either fully methylated or unmethylated. Approximately half of bivalent promoters with a high PDR in MM are associated with PC-PMDs. Therefore, the DNA methylation landscape of MM resembles that of the placenta, with stochastic methylated gain in CGIs embedded in large hypomethylated regions, suggesting that, as in other cancers, myeloma cells coopt placental nuclear programming [48, 49]; this finding is of particular interest since placental and cancerous tissues share relevant features such as immune modulation, angiogenesis and tissue invasion [50]. Interestingly, the specific enrichment of genes in immune-related pathways was revealed when we focused our analysis on genes located near PC-PMD boundaries.

PC-PMDs are also linked to the redistribution of repressive marks. We showed that PC-PMD hypomethylation is associated with a disruption in the H3K9me3 and H3K27me3 marks and variable depending on genomic regions, patients and DNA methylation levels. Promoter CGI controlling developmental genes are silenced by the PRC2 complex, which is responsible for the deposition of H3K27me3 via its catalytic component, EZH2. According to the model of Reddington et al [51], we can hypothesize that DNA methylation prevents PRC2 from binding to inappropriate targets, and that global hypomethylation due to tumorigenesis drives H3K27me3 redistribution responsible for the derepression of target genes and the repression of new genes. This redistribution may partially explain the derepression of *HOXA9* [52] and the de novo bivalent promoters [53] observed in MM. H3K27me3 redistribution could also play an important role in chromatin decompaction, as has been shown in ESCs [54], and could explain the increase in open chromatin regions within the heterochromatin state of MM samples [55]. These data highlight that PMDs must be considered as a separate entity in genomic analyses.

Furthermore, we showed that DNA hypomethylation is variable within the myeloma cells of one individual and that methylation loss modifies H3K27me3 distribution across patients; we can hypothesize that H3K27me3 redistribution is heterogeneous among myeloma cells, leading to cell-to-cell epigenetic variability. Consequently, the tumoral mass in a MM patient would be composed of an admixture of myeloma cells with divergent epigenetic identities, in accordance with what has been demonstrated recently in CLL [56].

## Conclusion

Altogether, our results show the importance of genome-wide epigenetic analysis and reveal marked PMD instability in MM patients at presentation. The perturbation in PMDs occurs at the epigenetic, transcriptomic and 3D organization levels and is responsible for interpatient variability. This disturbance in PMDs could explain, in part, some aberrant and heterogeneous phenotypes across MM samples [57, 58]. In addition, the lack of accurate global DNA methylation maintenance also drives intrapatient DNA methylation heterogeneity, which can contribute to intrapatient variability, allowing cell-to-cell diversity in transcriptional programs and opening multiple trajectories in response to therapy (Additional file 3: Figure S25).

## Materials and Methods

### Patients and samples

Myeloma PCs were derived from bone marrow samples collected at different stages of myeloma: 47 samples form 26 patients were collected at the SMM, diagnosis and relapse stages. Patients (15 women and 11 men, with a median age of 59 years) were monitored at the Intergroupe Francophone du Myélome (IFM) center, and all provided informed consent (Additional file 1: Table S1).

### Sample preparation and nucleic acid purification

Myeloma cells were purified using nanobeads and an anti-CD138 antibody (RoboSep, Stem Cell Technologies). After immunomagnetic sorting, the purity of the plasma cell suspension was verified, and only samples with at least 85% of PCs were subjected to genomic analysis. The average cell purity of MM was > 99% (range 90-100%). Control normal plasma cells (NPC1 to NPC3) were obtained from the bone marrow samples of three patients suspected of having monoclonal gammopathy of undetermined significance (MGUS), and the absence of abnormal PCs was assessed by flow cytometry. The average CD138+ cell purity was > 75%. The DNA and RNA of CD138+ cells were purified using the Qiagen protocol and the quality and quantity of the nucleic acids were measured on the Nanodrop, Qubit and Agilent profiles and stored in the MM biobanks of the Nantes and Toulouse Hospitals.

### eRRBS

eRRBS is an improvement of the reduced representation bisulfite sequencing (RRBS) protocol, resulting in an increase in CpG detection and coverage. eRRBS library preparations were performed by Integragen and adapted from the protocol described by Garrett-Bakelmanet al. [59]. DNA was digested with the Msp1 enzyme, fragments between 150 bp and 400 bp were selected, and bisulfite conversions were processed. Libraries were sequenced on a HiSeqTM 4000 Illumina machine using 75 bp paired-end reads to an average depth of 50X per covered CpG. The average number of reads sequenced per patient was 55 216 285; and the average alignment rate of uniquely mapped reads was 60.81%, with an average of 2 712 252 CpGs per patients with a coverage of 10X and an average of 1 538 510 CpGs per patient with a coverage of 60X (Additional file 4: Table S3).

### eRRBS analysis

The adaptor sequences were removed by Cutadapt (version 1.10) [60]. FastQC (version 0.11.4) was used for quality control of the Illumina paired-end sequencing data [61]. Bisulfite reads were aligned to the bisulfite-converted hg19 genome with the nondirectional model of Bismark alignment software (version 0.14.1) [62]. CpG methylation levels were obtained using the R package methylKit with the default settings: a minimum of 10 reads covering a CpG and at least 20 PHRED quality scores by CpG [63]. We calculated the epigenetic changes between two stages using methclone [17], an algorithm that detects loci of 4 adjacent CpGs (minimum depth of 60 reads), called epialleles (16 possible patterns according to CpG methylation).

### Epipolymorphism and epiallele shifting

Epipolymorphism (e) was calculated for each locus of the four adjacent CpGs, to measure intratumor epigenetic heterogeneity, as previously described by Landan et al. [18]:

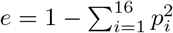

where *p*_*i*_ is the proportion of the epiallele i. The minimal epipolymorphism is 0 (only one pattern represented); epipolymorphism cannot have a value greater than 1. The maximum value of epipolymorphism is 0.9375 (when all 16 patterns are equally distributed). To quantify the degree of epiallele pattern shift, methclone was used to compute the entropy difference (ΔS) and thus compare the distributions of epialleles between different stages. The entropy difference value can range from 0 (no change) to −144 (maximum change). Loci are characterized as eloci when ΔS < −70 (corresponding to a significant entropy shift). The methclone algorithm allows the discovery, quantification and ranking of subclonal selection based on epiallele shifts. This allows us to measure clonal evolution between disease stages and epigenetic heterogeneity.

To normalize and compare the number of eloci per patient, we computed the number of eloci per million loci sequenced (EPM) as previously described by Sheng Li et al. [17]:

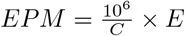

where E is the total number of eloci detected between the two stages and C is the total number of loci covered by both samples.

A locus between the NPC and diagnosis samples was defined as an elocus only if it was annotated as an elocus with the 3 NPCs. A hypermethylated or hypomethylated elocus was defined as a DNA methylation difference of at least 25%.

The PDR variable was defined as the percentage of epialleles exhibiting heterogeneous DNA methylation (i.e., not fully methylated or unmethylated).

### WGBS

Data quality control and adaptor trimmed were performed with the Trimmomatic tool [64]. Read mapping was carried out with the methylCtools aligner [31]. methylCtools uses an alignment approach similar to that of Bismark by improving the handling of large amounts of data and the speed of alignment. CpG extraction and methylation analyses were carried out with methylCtools and the bsseq R package [65], allowing the analysis, management and storage of WGBS data. As with eRRBS data, we set a coverage threshold of a minimum of 10X for each CpG sequenced for our analyses. The dmrseq R package was used to visualize WGBS data and smooth methylation signals [66].

### Detecting PMDs

PMDs were detected in WGBS samples using the R package MethylSeekR [28]. Prior to the detection of PMDs, CpGs overlapping with common single-nucleotide polymorphism (SNPs) were removed from the data (dbSNP137commonhg19, version 1.0.0). The distribution of *α* values was used to determine the presence or absence of PMDs in samples. When the *α* distribution was bimodal, the segmentPMDs function was run to identify PMDs on the genome by a hidden Markov model.

### RNA sequencing (RNA-seq)

RNA-seq was performed using 200 ng of total RNA by GATC Biotech. Directional libraries were generated after mRNA selection by polyA selection using the UTP method. RNA-seq libraries were sequenced on a HiSeq 2500 Illumina machine using 100 bp paired-end reads. The average number of reads sequenced per sample was 68 454 433. The average alignment rate of uniquely mapped reads was 66% (corresponding to 48 034 921 reads). Read alignment was performed using the STAR aligner (version 2.4.0f1) [67] and human genome hg19 as the reference. FastQC (version 0.11.4) [61] was used for basic quality control of the Illumina paired-end sequencing data. PCR duplicates were determined and removed using the Picard algorithm [68]. The number of reads mapped to each gene was calculated using HTSeq-count, part of the HTSeq framework [69], version 0.6.0. We then normalized the mapped read counts per million of mapped fragments (FPM) using the robust median ratio method with the DESeq2 R package [70].

### Hi-C data analysis and compartments A/B

Hi-C datasets were downloaded for three cell lines: GM12878 (GSM1608505), RPMI-8226 (GSM2334832) and U266 (GSM2334834). HiC-Pro software (version 2.10.0) [71] was used to process Hi-C data, from raw-data to normalized contact maps. All reads were mapped to hg19 using Bowtie2 (global parameters: ‒very-sensitive -L 30 ‒score-min L,−0.6,−0.2 ‒end-to-end ‒reorder; local parameters:–very-sensitive -L 20 ‒score-min L,−0.6,−0.2 ‒end-to-end ‒reorder). Contact maps were generated at 20 kb resolution and normalized by the iterative correction and eigen-vector decomposition (ICED) technique.

The 20-kb resolution intrachromosomal contact matrices generated by HiC-Pro were used as input to determine compartments A/B and to annotate and visualize interaction maps with R package HiTC version 1.22.1 [72]. Principal component analysis was used to separate chromatin into two compartments: compartment A, with higher gene density, and compartment B, with lower gene density. The determination of compartments A and B was estimated by the analysis of the eigenvectors of the genome contact matrix by the observed-expected method. On the basis that changes in the sign of the eigenvector of the contact matrix correspond to the limits of the genome compartments and taking into account gene density, the compartmentalization of the genome was defined.

### TADs

Hi-C data were used to determine the TADs with the HiCExplorer tool, a set of programs used to process, normalize, analyze and visualize Hi-C data [73]. TADs were defined with hicFindTADs and hicPlotTADs was used to visualize the TADs. For the comparison of PMD and TAD borders, we generated a set of randomized PMDs with a size and genomic distributions similar to those of real PMDs. We then calculated the distance between TAD borders and, on the one hand, PMD borders, and, on the other hand, random PMD borders. A t test was carried out to compare the distributions of these distances.

### Genomic annotation

RefSeq annotation and CpG islands were obtained from UCSC (https://garance/genome.ucsc.edu/) using the February 2009 (GRCh37/hg19) assembly. CpG shores were defined as regions flanking 2 kb of CpG islands, and CpG shelves were defined as regions flanking 2 kb of CpGs shores.

HOMER was used to annotate loci and eloci, using the annotatePeaks.pl script [74], which determines the genomic type annotation occupied by the center of the loci. We used their basic annotation, including TSSs, transcription termination sites (TTSs), exons, introns and intergenic regions. We defined the promoters as regions flanking 2 kb of the TSS, and we defined promoter CGIs as promoters that intersect with CpG islands.

All 127 reference epigenomes with 25 chromatin state segmentation annotations were downloaded from the NIH Roadmap Epigenomics Project (https://egg2.wustl.edu/roadmap/data/byFileType/chromhmmSegmentations/ChmmModels/imputed12marks/jointModel/final/;all\OT1\textquotedblleft25_imputed12marks_mnemonics.bed.gz\OT1\textquotedblrightfiles).

To measure the intersections between regions, bedtools was used (more precisely, the “intersect” option) [75].

Data from the BLUEPRINT Consortium were converted to hg19 coordinates using the liftOver tool [76].

### Functional annotation

GREAT tools (version 3.0.0) was used to assign biological functions to noncoding genomic regions by analyzing the annotations of the nearby genes [77] under the following default gene regulatory domain definitions; basal promoter: 5 kb upstream, 1 kb downstream and extension up to 1 Mb. Enrichment statistics were computed using the binomial test and the hypergeometric gene-based test. Pathways were selected as significantly enriched if the false discovery rate (FDR q-value) was < 0.01.

The Database for Annotation, Visualization and Integrated Discovery (DAVID) web interface (version 6.7) was used to perform functional enrichment analysis from a list of genes, especially the KEGG pathways functional database [78]. Enrichment statistics were computed using the Fisher test. For the significance threshold, we have considered genes as greatly enriched if the annotation categories yielded a p-value less than 0.1 (DAVID default threshold)[79].

### Survival analysis

Time to relapse data were available for 17 patients. For the relapse-free survival (RFS) analysis, the survival endpoint in this study was the time from diagnosis until relapse. The patients were divided into two groups by the median EPM value: low and high EPMs. Survival curves were estimated using the Kaplan-Meier method and compared with the log-rank test.

For all statistical tests in this study, a two-sided p-value of 0.05 was considered statistically significant. All statistical analyses were performed with software R 3.5.1 [80], in addition to the packages already mentioned in the text, packages data.table [81], viridis, ggbio, GenomicRanges, RColorBrewer, biovizBase, grid, gridBase, MASS, fields, KernSmooth, sp, org.Hs.eg.db, DBI and survival were used.

## Abbreviation

MM: Multiple myeloma
PMDs: Partially methylated domains
CGI: CpG island
TSS: Transcription start site
PCs: Plasma cells
NPCs: Normal plasma cells
WGBS: Whole genome bisulfite sequencing
eRRBS: Enhanced reduced representation bisulfite sequencing
SMM: Smoldering multiple myeloma
PDR: Proportion of discordant reads
eloci: Epigenetic loci
EPM: Eloci per million loci covered
ESCs: embryonic stem cells
FPM: mapped read counts per million of mapped fragments
OR: Odd ratio
CLL: Chronic lymphocytic leukemia
TAD: Topological associated domain
LAD: lamina-associated domain
KEGG: Kyoto encyclopedia of genes and genomes
3D: Three-dimensional
IFM: Intergroupe francophone du myélome
MGUS: Gammopathy of undetermined significance
RRBS: Reduced representation bisulfite sequencing
SNPs: Single-nucleotide polymorphism
RNA-seq: RNA sequencing
ICED: Iterative correction and eigenvector decomposition
TTS: Transcription termination site
FDR: False discovery rate
DAVID: Database for annotation, visualization and integrated discovery
RFS: Relapse-free survival

## Acknowledgements

This study makes use of data generated by the BLUEPRINT Consortium. A full list of the investigators who contributed to the generation of the data is available from www.blueprint-epigenome.eu. Funding for the project was provided by the European Union’s Seventh Framework Programme (FP7/2007-2013) under grant agreement no 282510 – BLUEPRINT.

## Author’s contributions

S.M. and F.M. designed the study, collected, and analyzed the data and wrote the paper; J.D. and C.H. collected and analyzed the data and wrote the paper; M.D, E.D, N.R, H.A.-L., O.D., T.F., J-P. M., R.E., M.P. and N.M. collected the data; C.G.-C., V.G. and L.C. analyzed the data.

## Availability of data

WGBS, eRRBS and RNA-seq data have been submitted to the European Genome-phenome Archive (https://www.ebi.ac.uk/ega) under dataset accession EGAS00001004346, EGAS00001004348, EGAS00001004347, respectively.

WGBS Blueprint data of normal plasma cells and multiple myeloma patient are available from the EGA under the accessions EGAD00001002322 and EGAD00001002521 respectively. ChIP-Seq data of multiple myeloma patient are available from the EGA under the accessions EGAD00001002379.

## Ethics approval and consent to participate

All the samples were obtained from patients enrolled in the different randomized clinical trials implemented by the Intergroupe Francophone du Myelome (IFM); IFM 99-02, IFM 99-06, IFM 2005-01, FIRST/IFM 2007-01, IFM 2007-02, Genomgus Study and IFM 2009 (www.clinicaltrials.gov) or routine practice in France. Samples and data were obtained after an informed consent was signed. All research procedures conformed to the principles of Helsinki Declaration.

## Consent for publication

Not applicable.

## Additional Files

Additional file 1 — Table S1

Data resource.

Additional file 2 — Table S2

Distribution of CpG sites within 100 bp genomic units in the human genome.

Additional file 3 — Figures S1–S25

Supplementary figures.

Additional file 4 — Table S3

Description of sequencing metrics for eRRBS data.

Additional file 5 — Table S4

Average number of CpGs covered by eRRBS with read depths greater than 10X and 60X, given by genomic features.

Additional file 6 — Table S5

Odds ratios and 95% confidence intervals for association between gene expression and promoter methylation according to low or high PDR levels in the promoter.

