## Supplementary Figures for "The DNA methylation landscape of multiple myeloma shows extensive inter- and intrapatient heterogeneity that fuels transcriptomic variability"

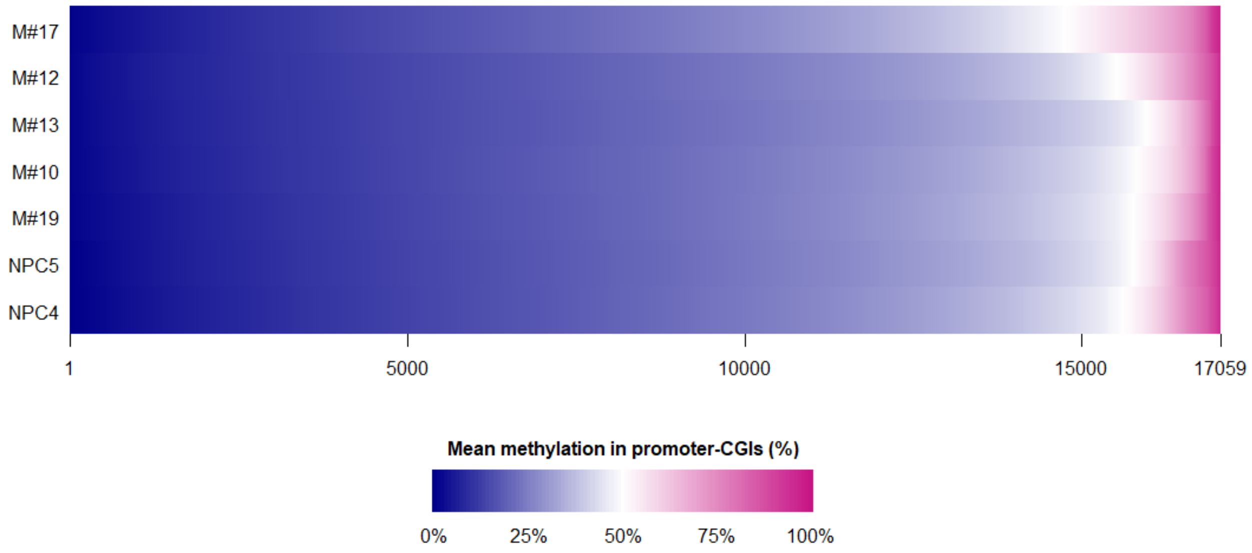


**Figure S1 |** Mean methylation levels in promoter-CGIs across MM patients and NPCs.

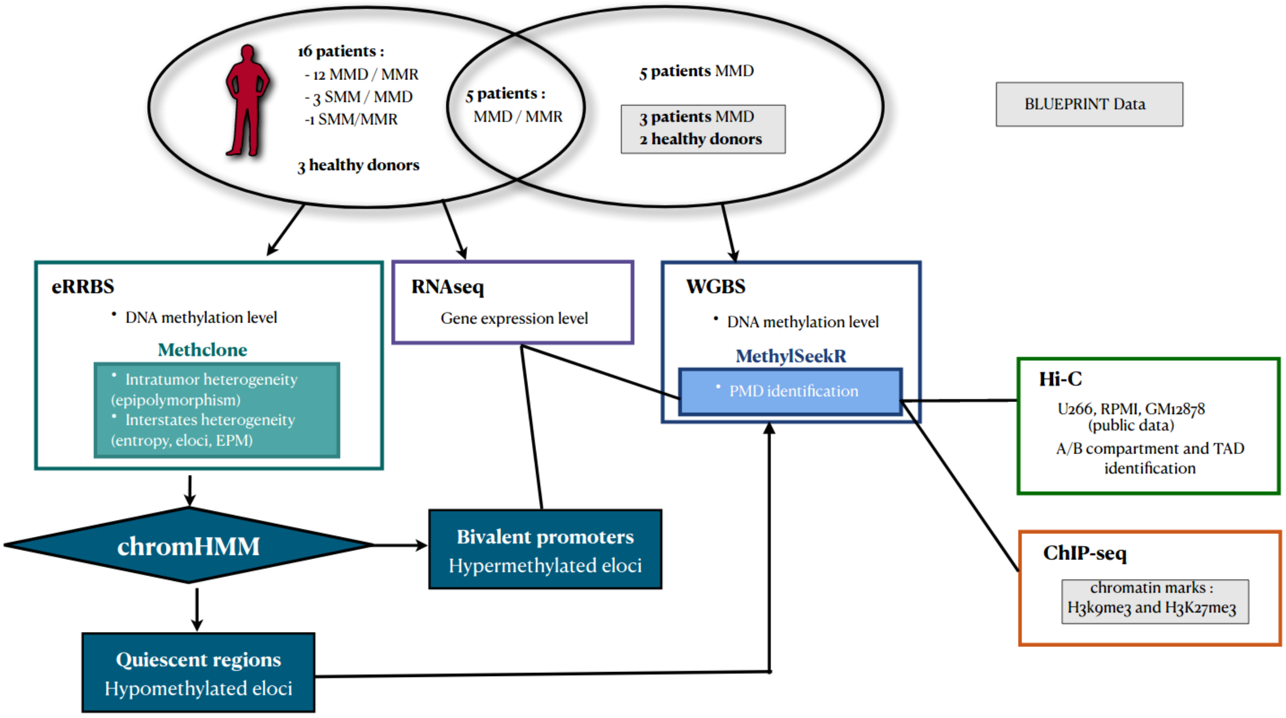


**Figure S2 |** Diagram of the analytical process.

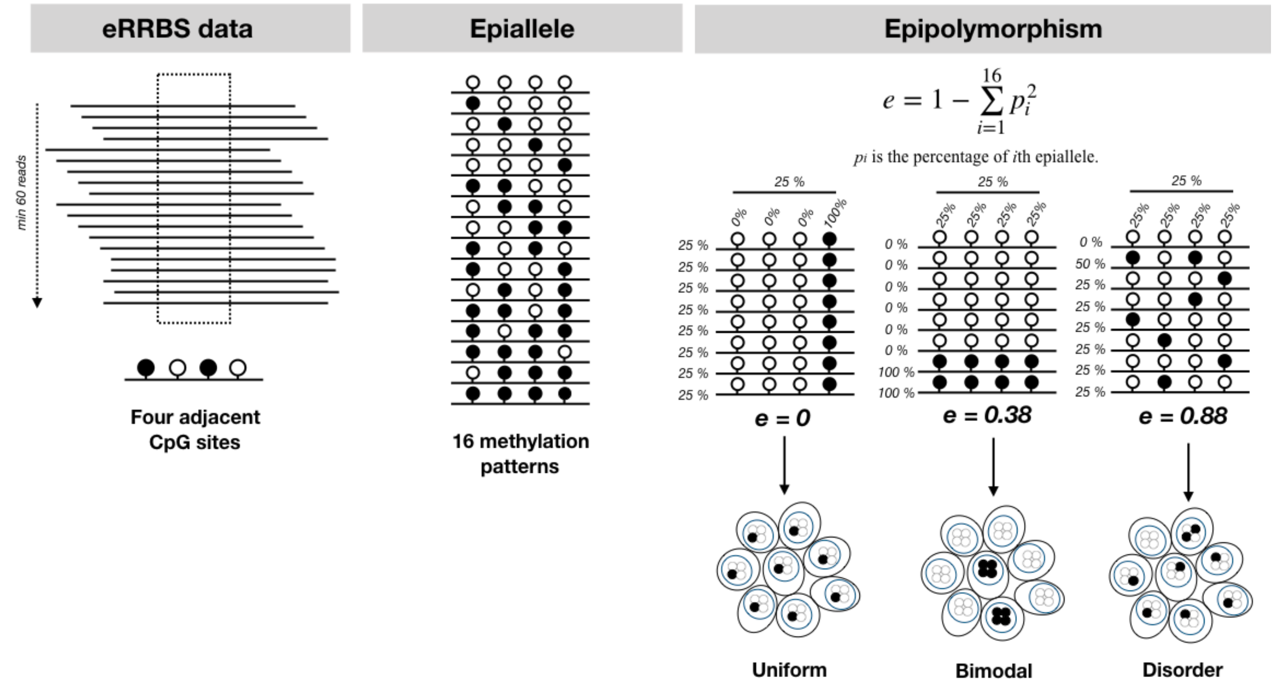


**Figure S3 |** Loci detection of 4 adjacent CpGs and measurement of their epipolymorphism with the methclone tool. Identification of regions containing 4 adjacent CpGs at least 60 reads covered (= loci; filled black circle: methylated CpG; empty circle: unmethylated CpG). Sixteen methylation patterns of the epiallele from 4 adjacent CpGs are possible. Assessment of the epipolymorphism value (i.e., the degree of methylation heterogeneity) of each locus, taking into account the distribution of the probabilities of these 16 possible methylation patterns. For the same average level of methylation, different epipolymorphism values are possible depending on the bulk.

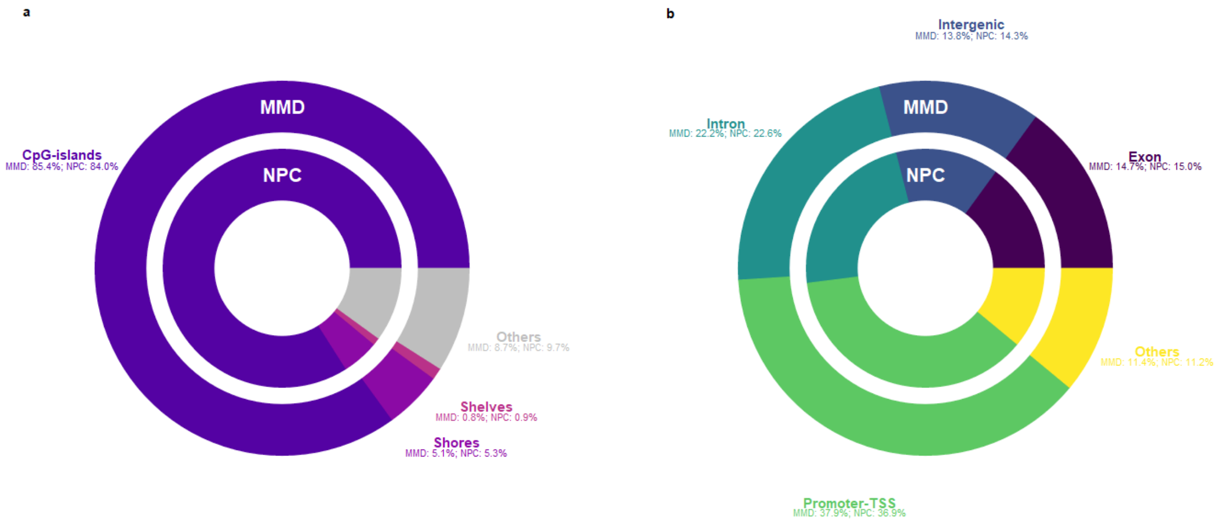


**Figure S4 |** Genome distribution of background loci across MM patients and NPCs in CpG islands, shores and shelves (a), and in promoter TSS, intron, exon and intergenic regions (b).

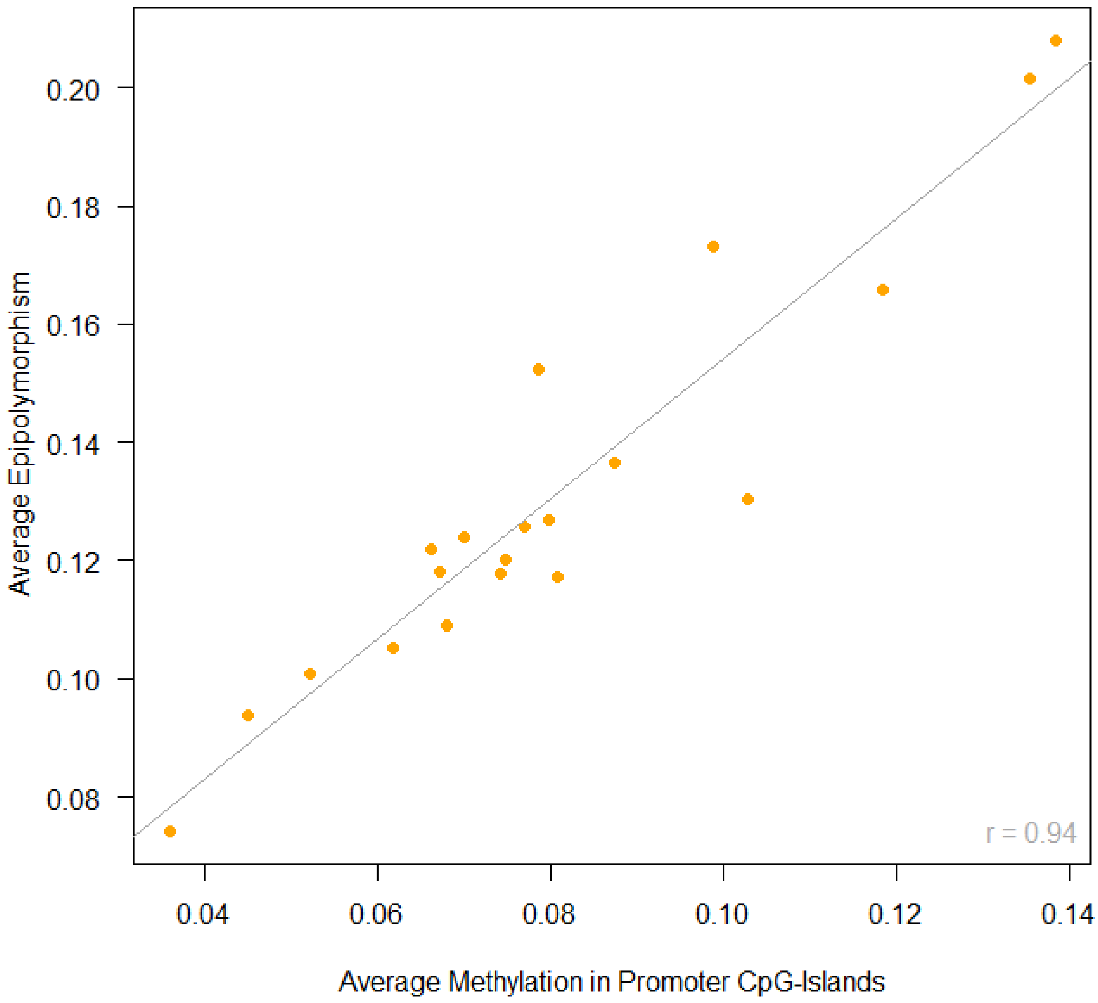


**Figure S5 |** Correlation between average promoter CGI methylation and epipolymorphism at diagnosis per sample.

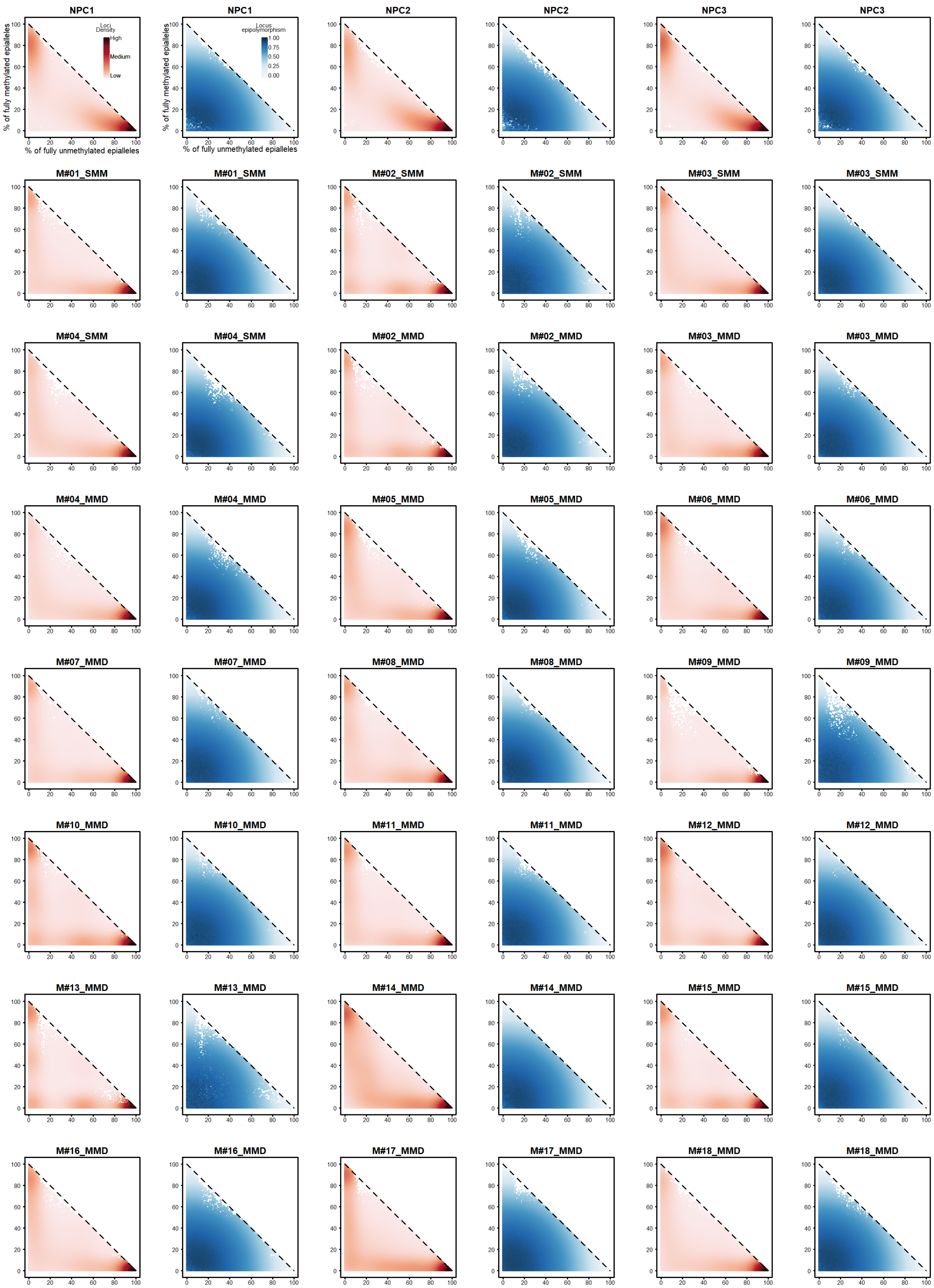
NPC1
NPC1
NPC2
NPC2
NPC3
NPC3
% of fully methylated epialleles
% of fully unmethylated epialleles
Loci Density
High
Medium
Low
Locus epipolymorphism
1.00
0.75
0.50
0.25
0.00
0 20 40 60 80 100
M#01_SMM
M#01_SMM
M#02_SMM
M#02_SMM
M#03_SMM
M#03_SMM
M#04_SMM
M#04_SMM
M#02_MMD
M#02_MMD
M#03_MMD
M#03_MMD
M#04_MMD
M#04_MMD
M#05_MMD
M#05_MMD
M#06_MMD
M#06_MMD
M#07_MMD
M#07_MMD
M#08_MMD
M#08_MMD
M#09_MMD
M#09_MMD
M#10_MMD
M#10_MMD
M#11_MMD
M#11_MMD
M#12_MMD
M#12_MMD
M#13_MMD
M#13_MMD
M#14_MMD
M#14_MMD
M#15_MMD
M#15_MMD
M#16_MMD
M#16_MMD
M#17_MMD
M#17_MMD
M#18_MMD
M#18_MMD

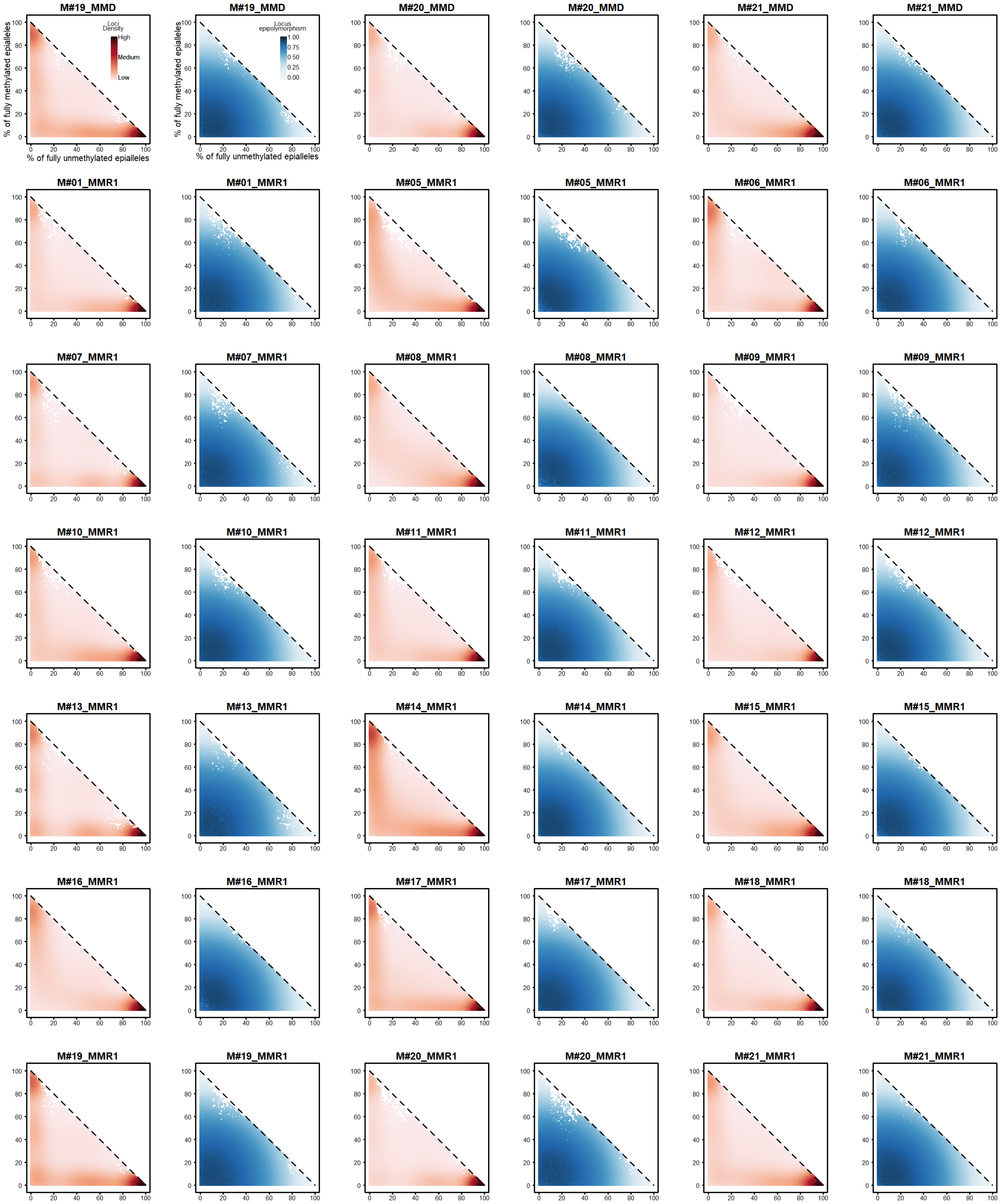


**Figure S6 |** Scatterplots showing loci organization as in Figure 2b for NPCs and all paired samples (SMM/Diagnosis/Relapse). Each point corresponds to a locus of 4 adjacent CpGs. Shown on the left is a representation where each point is color coded according to the density of the surrounding points; on the right each point is color coded according to its level of epipolymorphism.

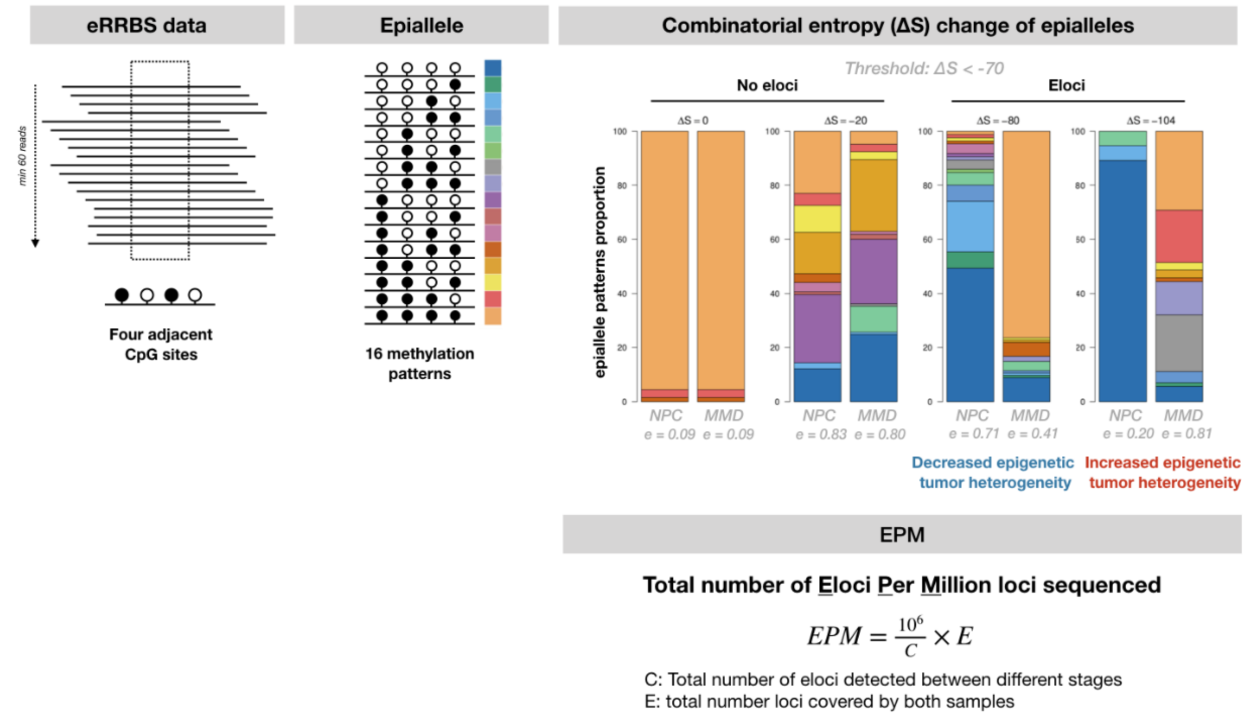


**Figure S7 |** Detection of epiallelic changes. From a combinatorial entropy value (threshold: ΔS <-70), Methclone determines loci with a significant change in epiallelic composition (= eloci). Filled black circle: methylated CpG; empty circle: unmethylated CpG.

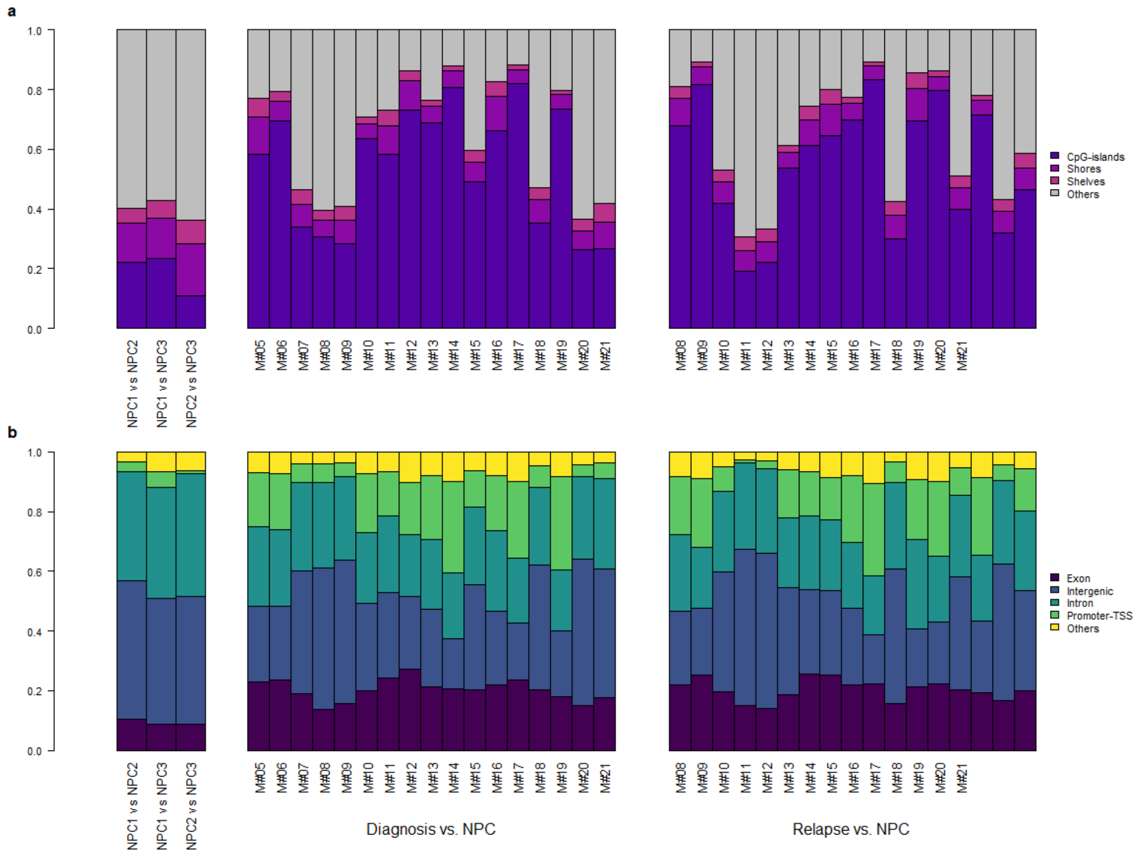


**Figure S8 |** Genomic distribution of NPCs versus NPCs eloci (to the left), NPCs versus diagnosis eloci (at the middle) and NPCs versus relapse eloci (to the right); in CGIs and adjacent regions (a) and in the main genomic regions: promoter TSS, exons, introns and intergenic regions (b). The distribution of NPCs versus NPCs eloci is shown as a control.

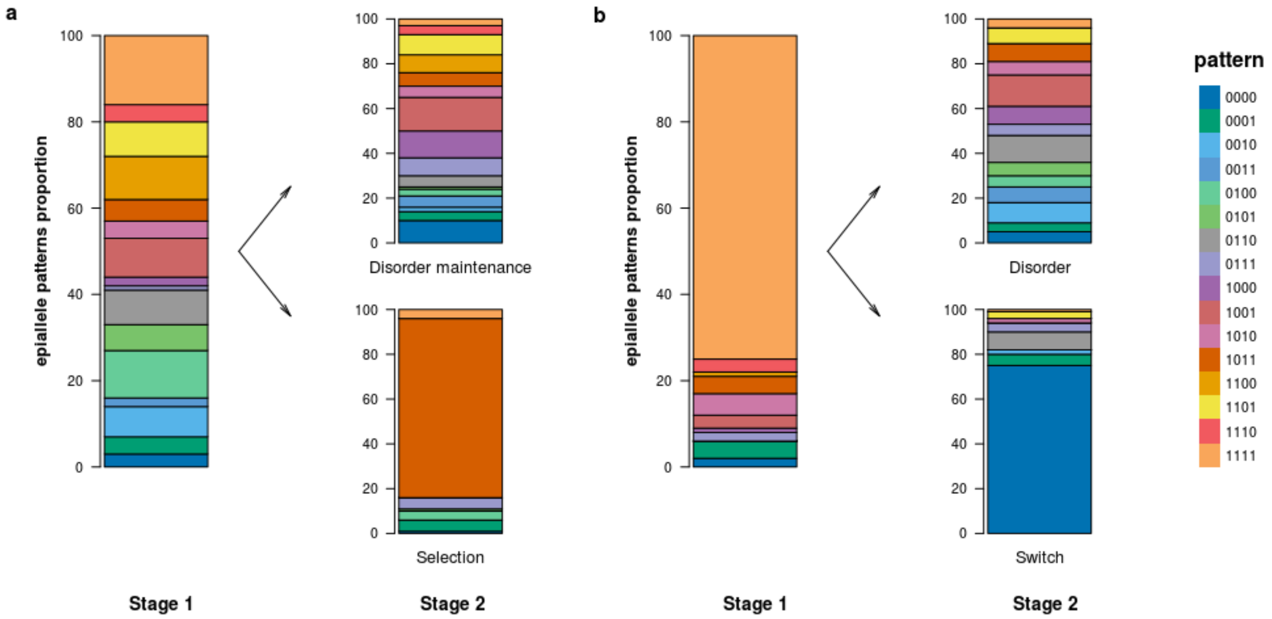


**Figure S9 |** Illustration of the four extreme changes in epiallele patterns between two stages: (a) Stage 1, heterogeneous epialleles; stage 2 (top), a similar heterogeneous pattern; stage 2 (bottom), selection of one pattern. (b) Stage 1, major fully methylated pattern; stage 2 (top), heterogeneous pattern; stage 2 (bottom), major fully unmethylated pattern. Legend for methylation patterns: 1 = methylated CpG and 0 = unmethylated CpG.

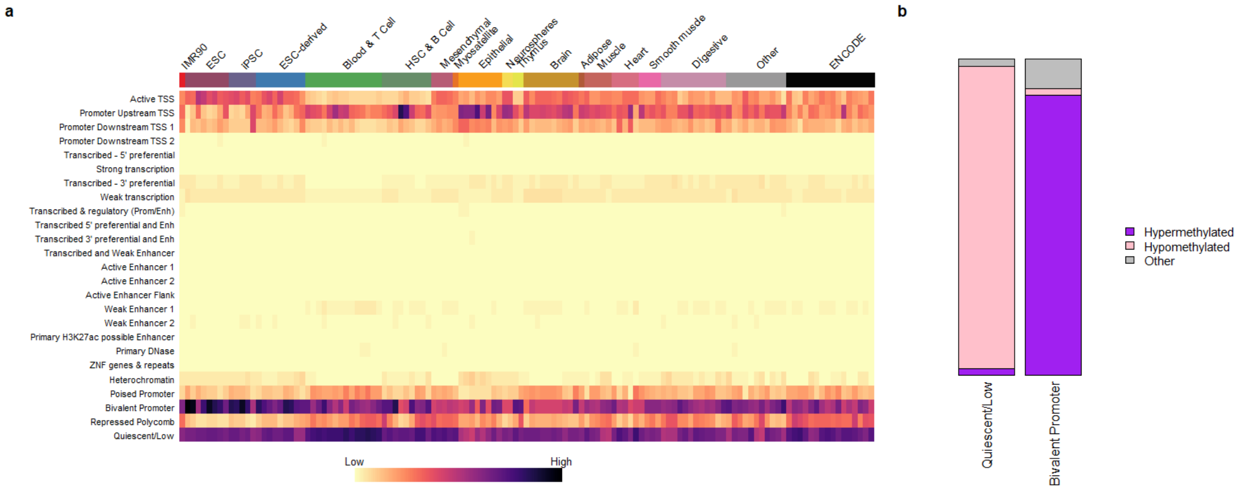


**Figure S10 |** Eloci (NPC vs diagnosis) annotation according to the chromatin states defined by ChromHMM. (a) Eloci annotation between NPC and MM at diagnosis samples according to the 25 chromatin states defined by ChromHMM for 127 epigenomes available on UCSC. (b) Proportion of hypo- and hypermethylated eloci in bivalent promoter and quiescent regions defined by ChromHMM.

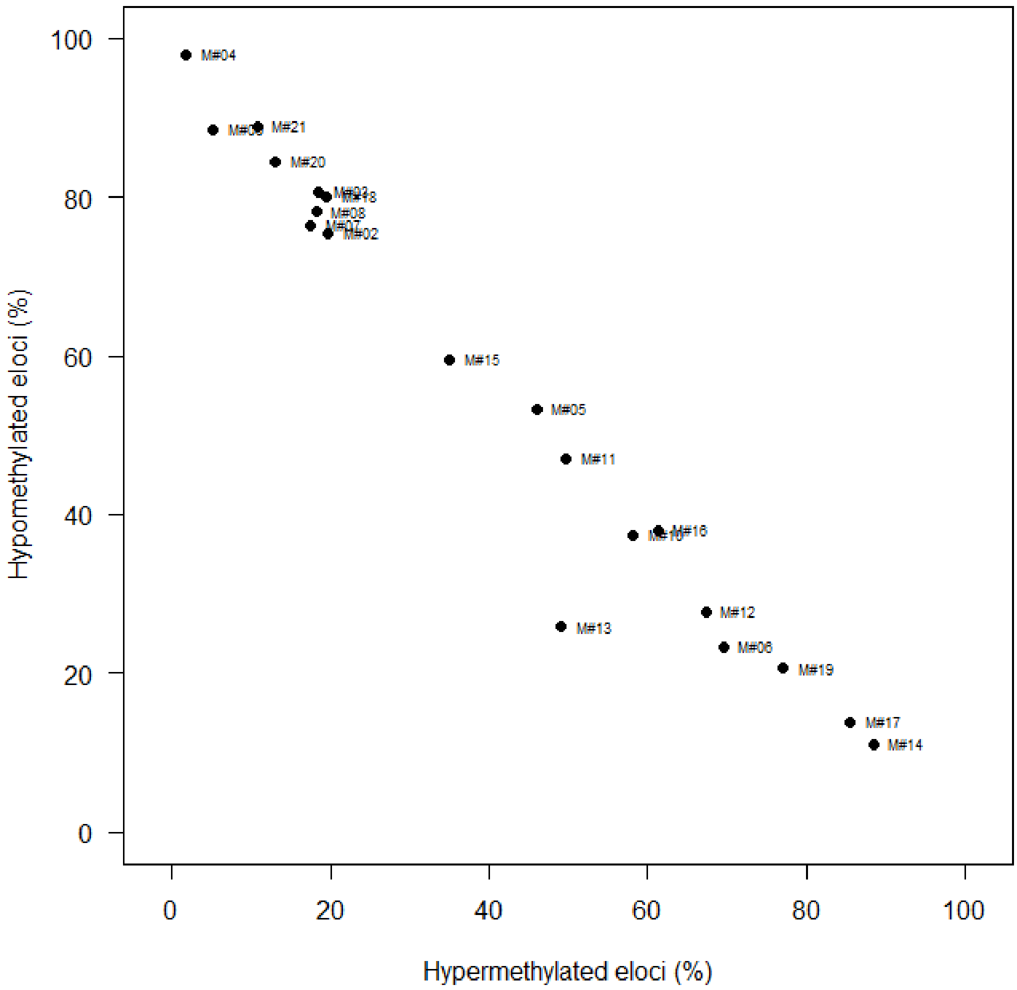


**Figure S11 |** Distribution of MM samples at diagnosis according to their percentage of hypo- and hypermethylated eloci (NPC vs diagnosis).

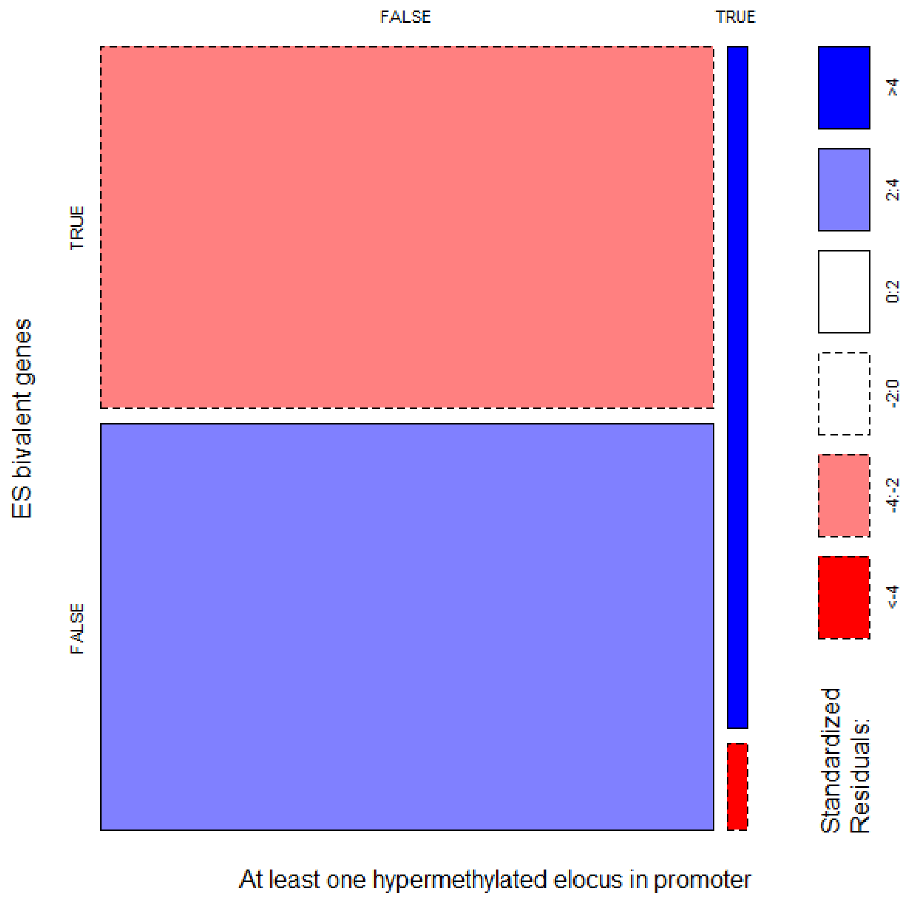


**Figure S12 |** Enrichment of hypermethylated eloci in the bivalent promoters of embryonic stem cells.

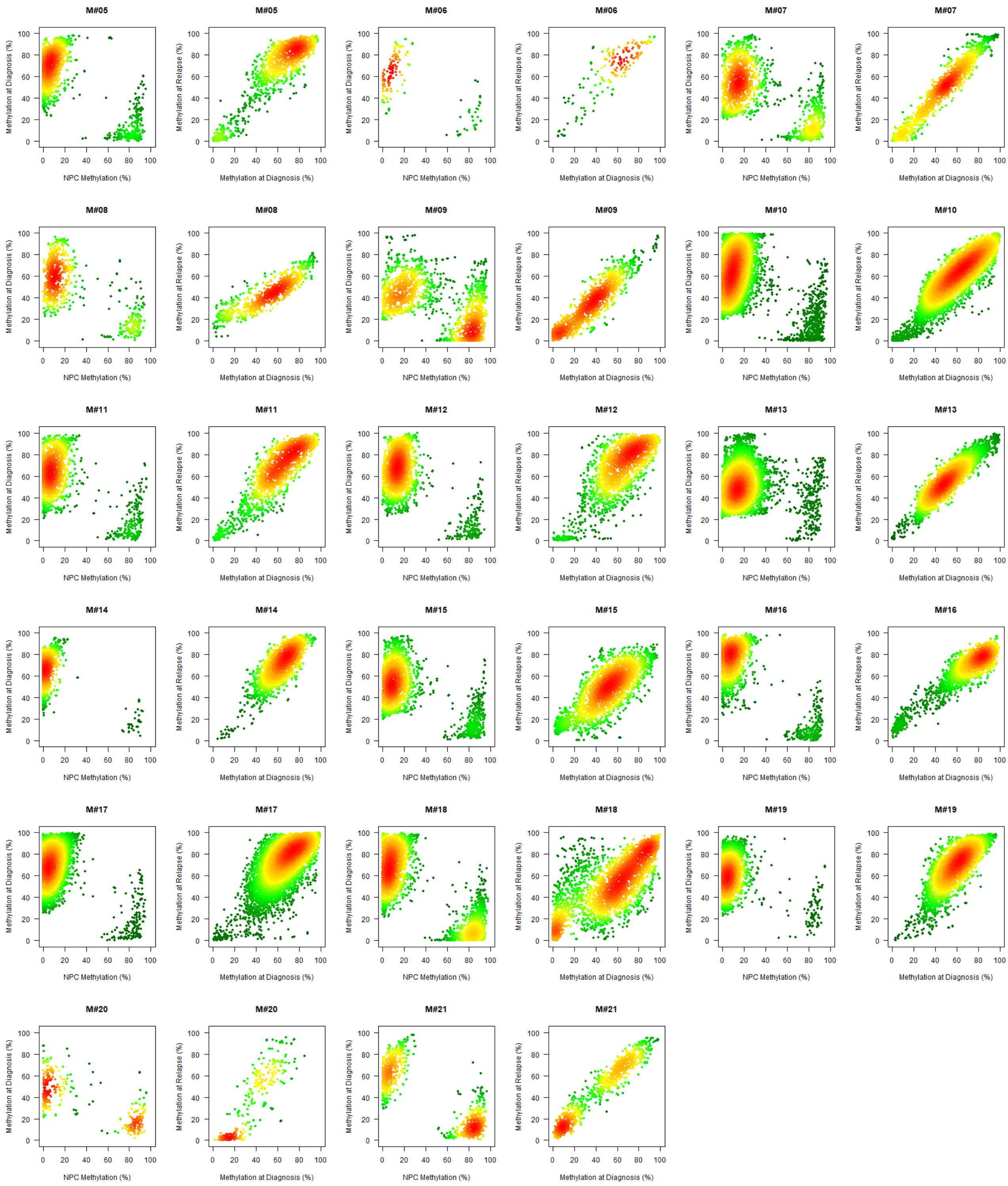


**Figure S13 |** Methylation disruption in bivalent promoters in MM. Scatterplot of eloci in bivalent promoters as a function of DNA methylation in NPCs and diagnosis samples (left figure) and as a function of DNA methylation at diagnosis and relapse (right figure) for all patients. The color gradient corresponds to the point density (low is green; high is red).

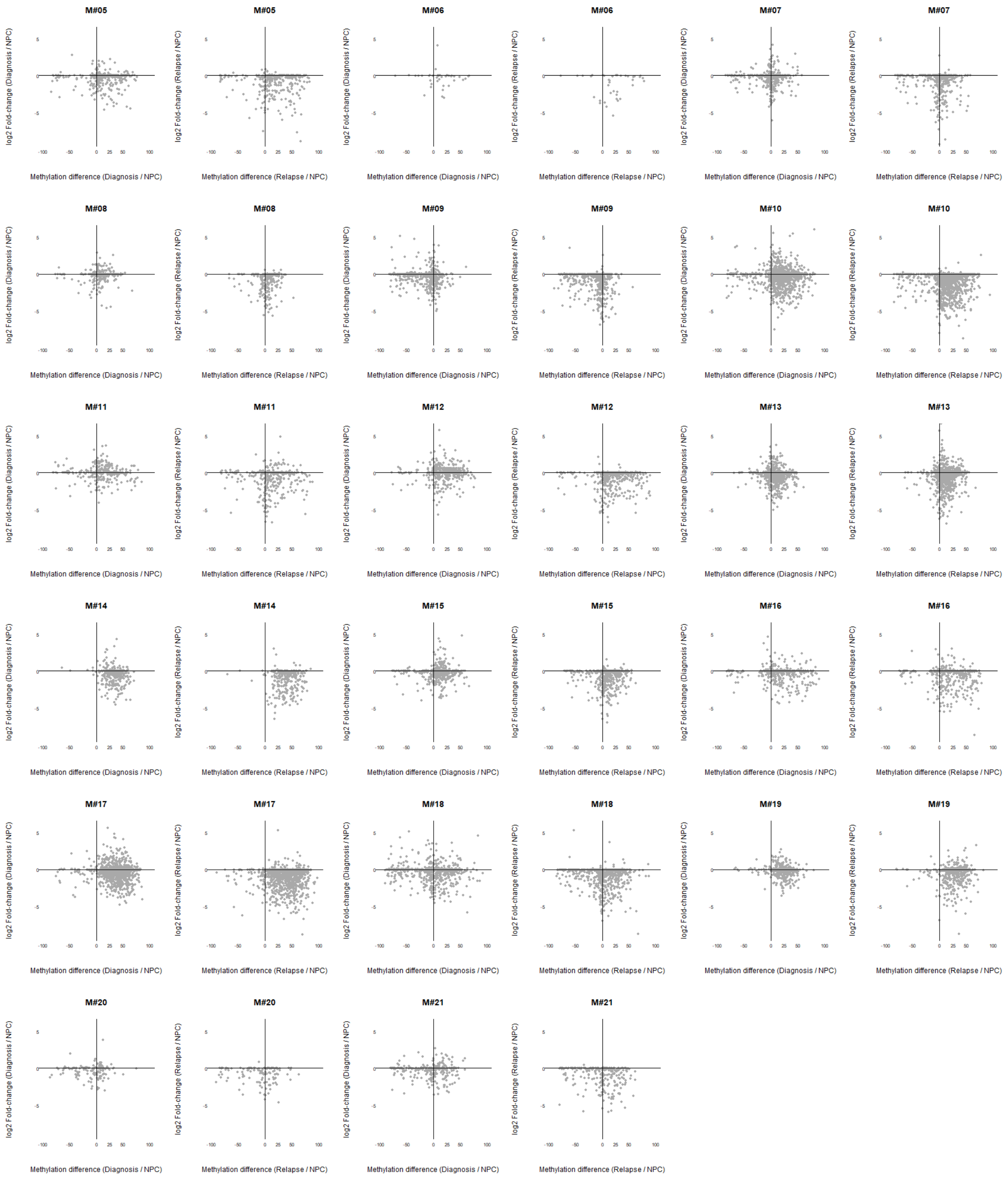


**Figure S14 |** Expression evolution of genes with eloci in bivalent promoters. Scatterplot of genes depending on the methylation level in the promoter and expression levels (left: diagnosis/NPC; right: relapse/NPC).

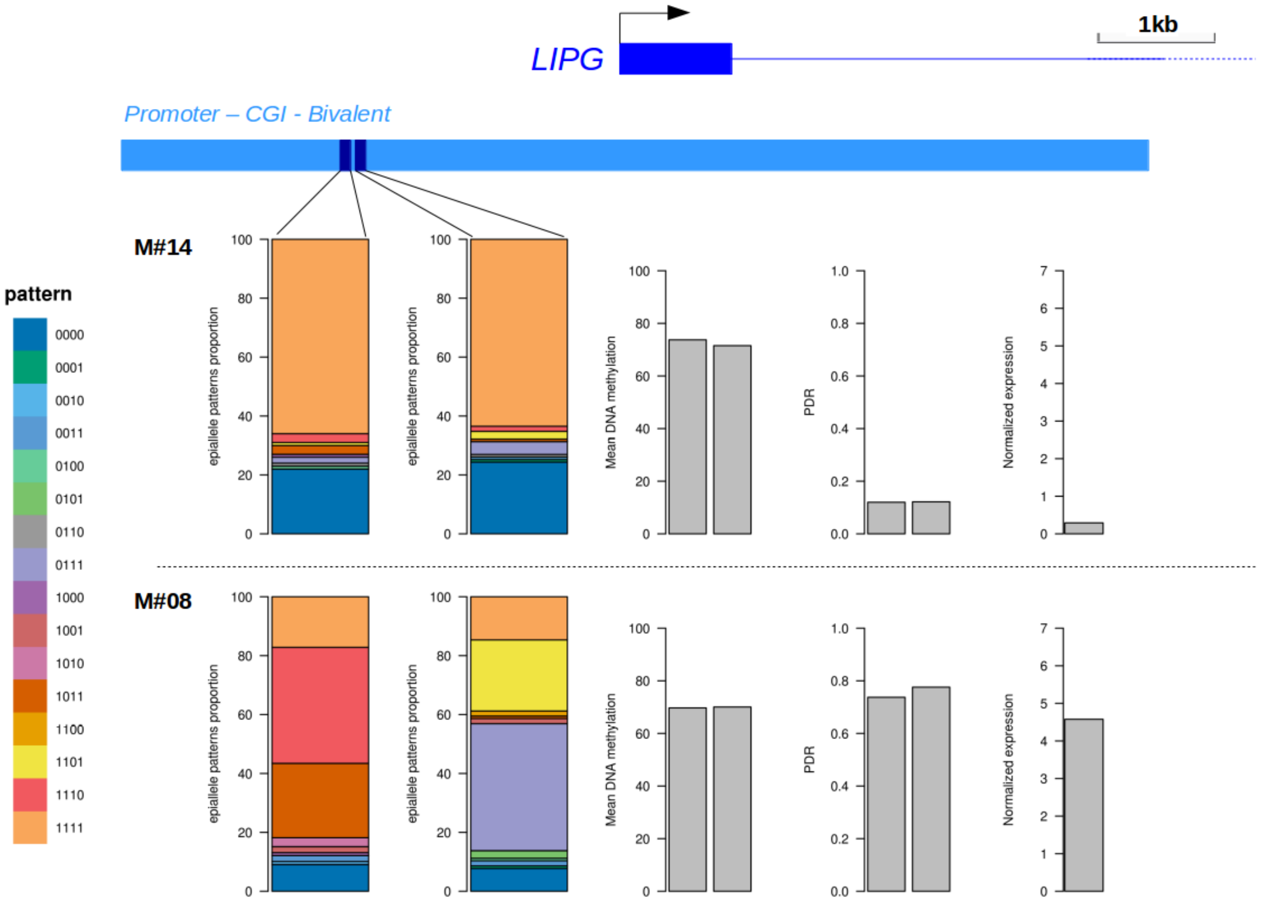


**Figure S15 |** Example of the decoupling relationship between promoter methylation and gene expression. The promoter region of *LIPG* shows comparable methylation levels in two samples (M#14 and M#08) but different PDR and expression levels (right). Epiallele patterns of two loci are shown in the illustration (left). Legend for methylation patterns: 1 = methylated CpG and 0 = unmethylated CpG.

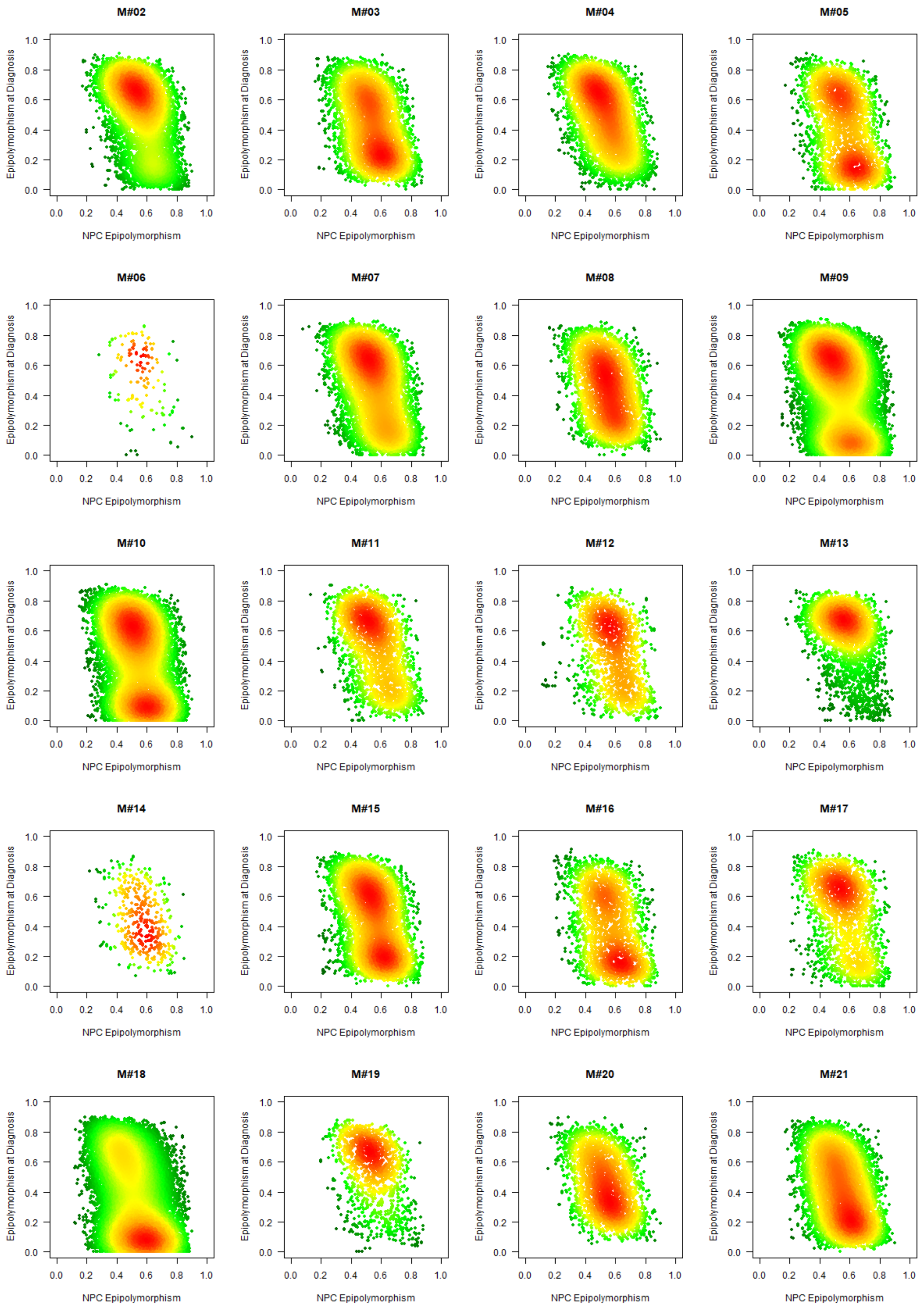


**Figure S16 |** Characterization of hypomethylated eloci epipolymorphism evolution. Scatterplot of eloci as a function of their epipolymorphism in NPC and diagnosis samples for all patients. The color gradient corresponds to the point density (low is green; high is red).

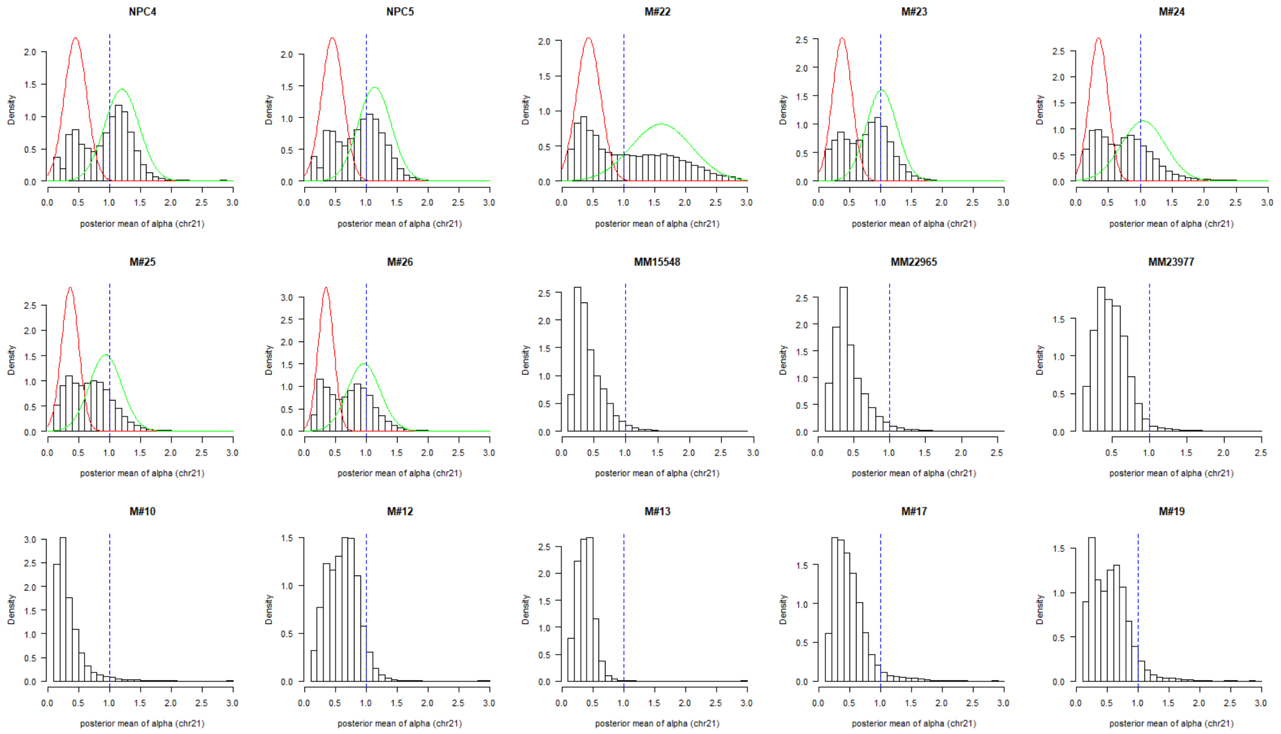


**Figure S17 |** Distribution of the α-value for chromosome 21. This distribution is similar for all chromosomes. The α-value characterizes the distribution of methylation levels in genomic windows of 100 CpGs. If the distribution of α-values is bimodal or has a large fraction of α value greater than or equal to 1 (blue dotted line), then this is evidence of the presence of PMDs. In the presence of a bimodal distribution, it is possible to calculate the PMDs on the genome with a hidden Markov model, whose two adjusted Gaussian distributions are represented in green and red.

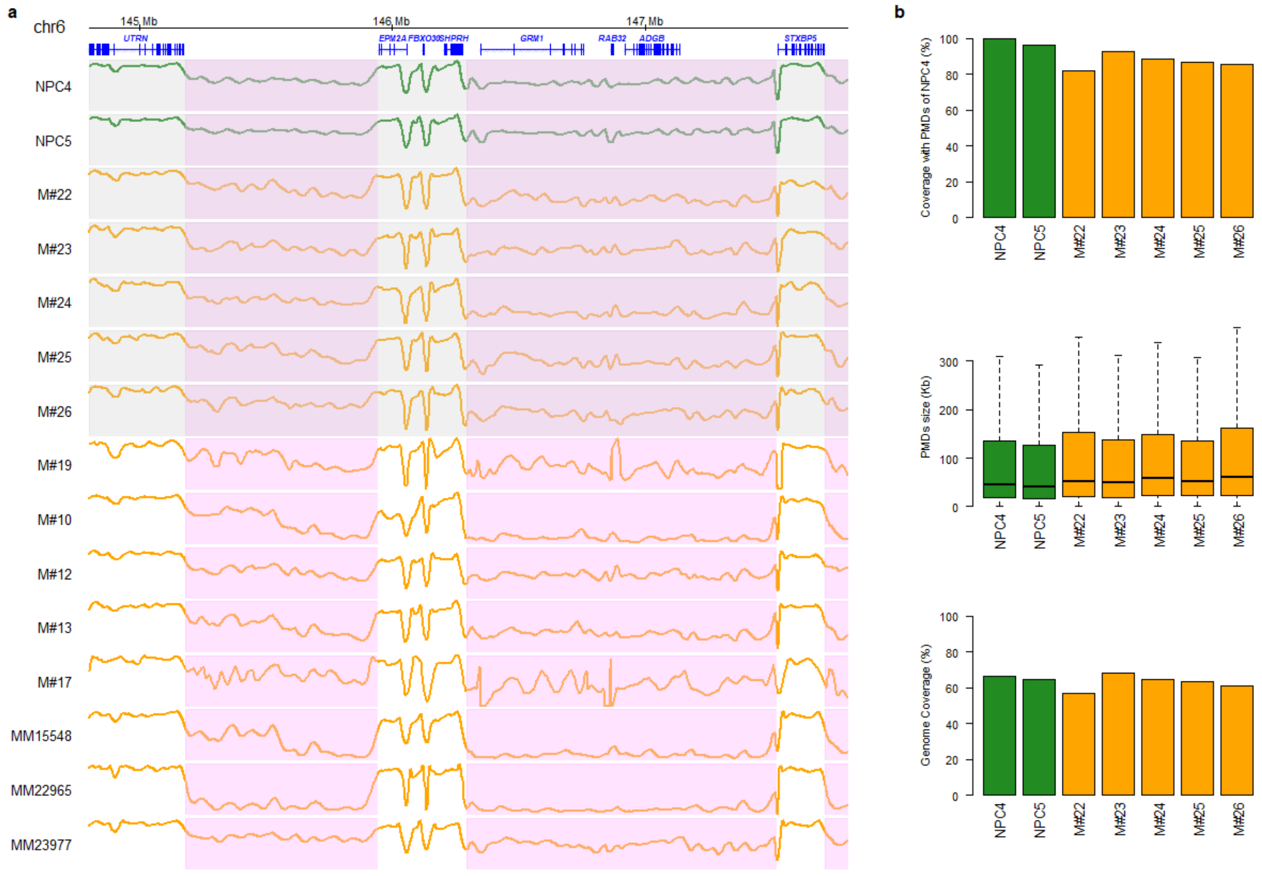


**Figure S18 |** Comparison of NPC and MM PMDs. (a) WGBS DNA methylation visualization in a genomic region encompassing two PMDs (pink area) in sample NPC4. Green lanes correspond to NPC methylation profiles, and orange lanes correspond to MM methylation profiles at diagnosis. The shaded profiles correspond to samples in which PMDs could be detected. (b) General characteristics of PMDs.

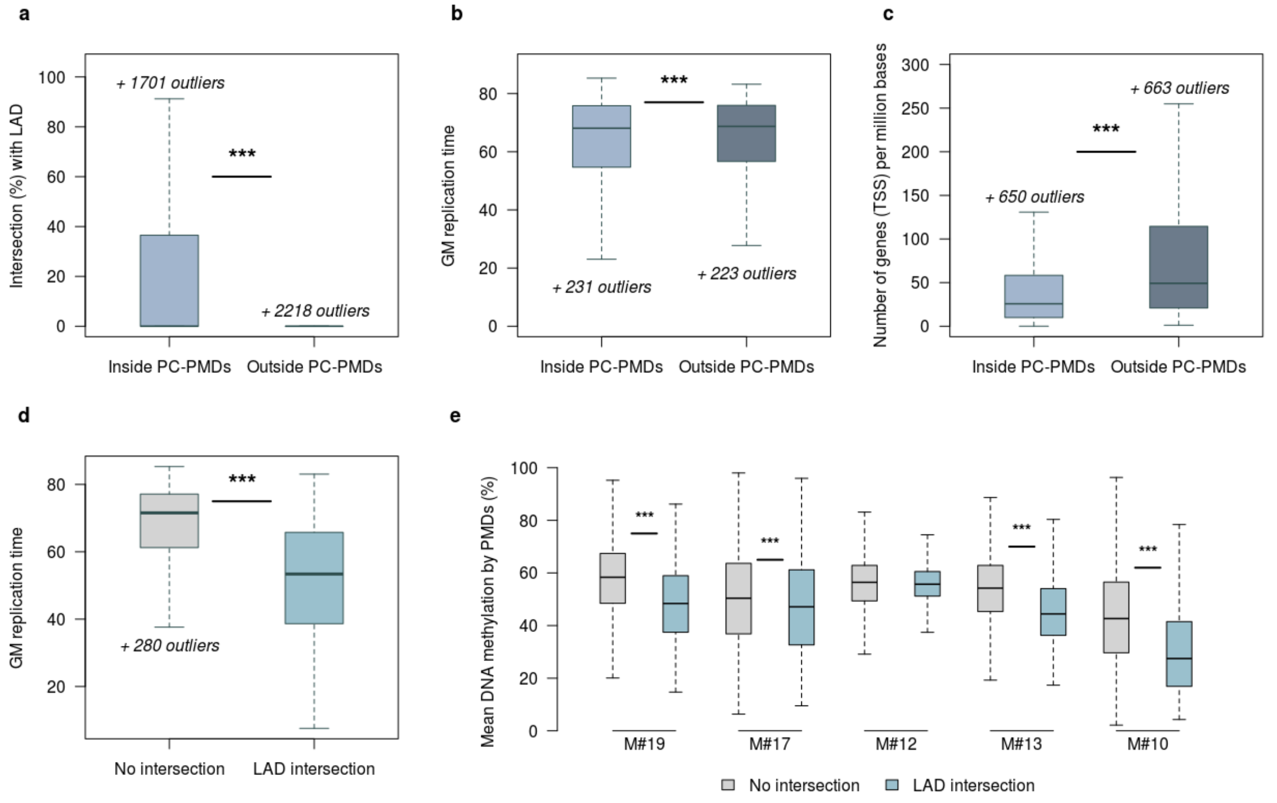


**Figure S19 |** Characteristics of PC-PMDs. (a) Percentage of intersections between LADs [30] and regions inside and outside PC-PMDs. (b) Replication timing of the GM12878 cell line (ENCODE data) inside and outside PC-PMDs. (c) Number of genes per million bases inside and outside PC-PMDs. (d) Replication timing of the GM12878 cell line in PC-PMDs intersecting (or not) with an LAD. (e) Mean DNA methylation level in PC-PMDs intersecting (or not) with an LAD per patient.

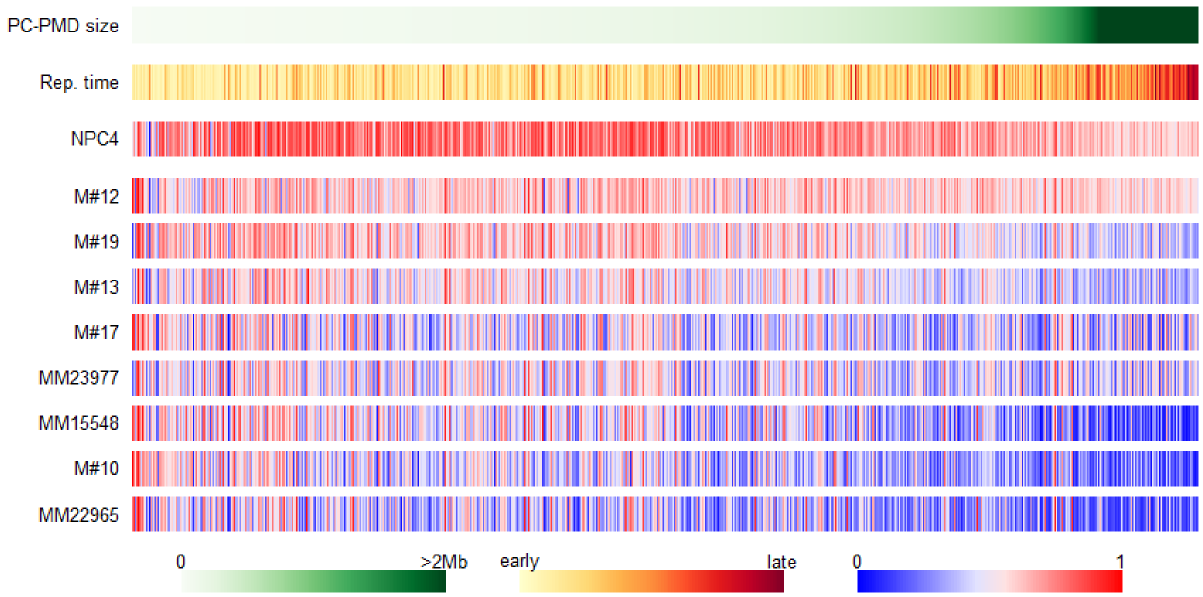


**Figure S20 |** Heatmap of PC-PMDs, sorted by increasing size, associated with GM12878 replication timing (ENCODE data) and DNA methylation levels in WGBS samples (NPC and diagnosis).

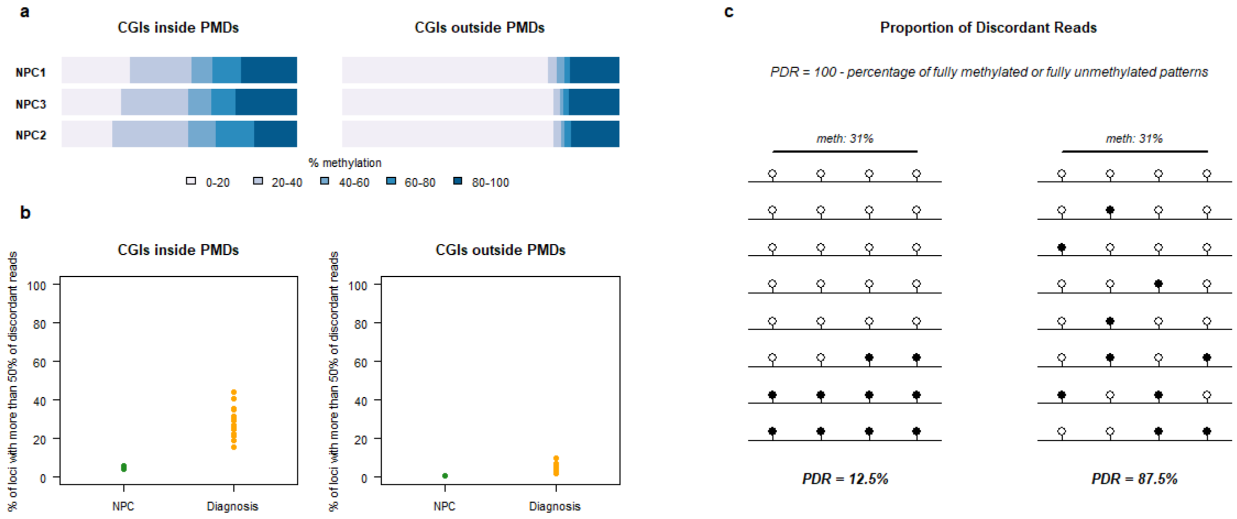


**Figure S21 |** Homogeneous CGI methylation perturbation in NPCs (a) Proportion of CGIs with partial methylation inside and outside PC-PMDs (b) Proportion of loci with more than 50% discordant reads inside and outside PC-PMDs. (c) PDR definition; for the same average level of methylation, different PDR values are possible.

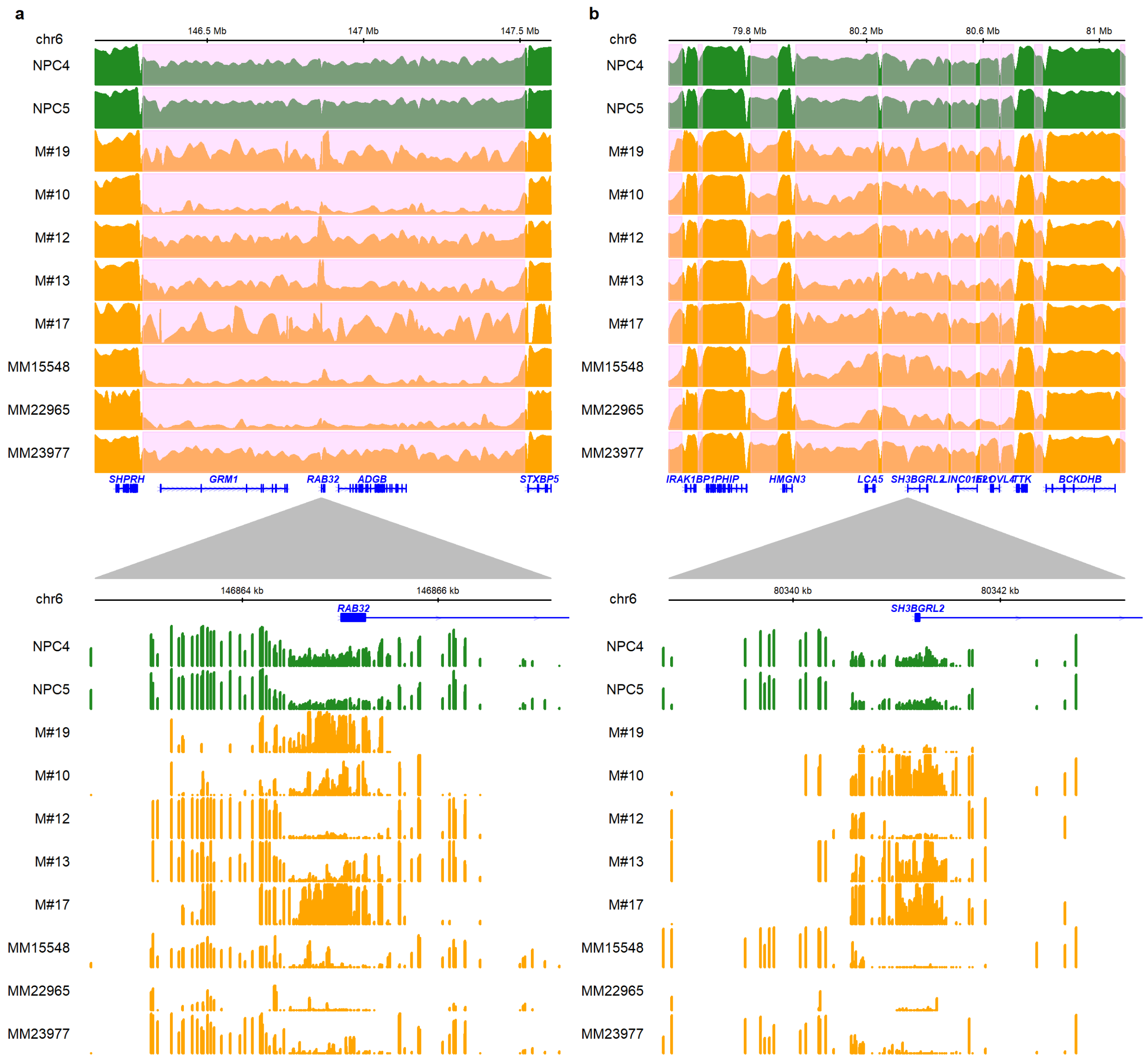


**Figure S22 |** Examples of genes with bivalent promoter CGIs with disrupted DNA methylation regions embedded within large partially methylated regions (pink area): *RAB32* (a) and *SH3BGRL2* (b).

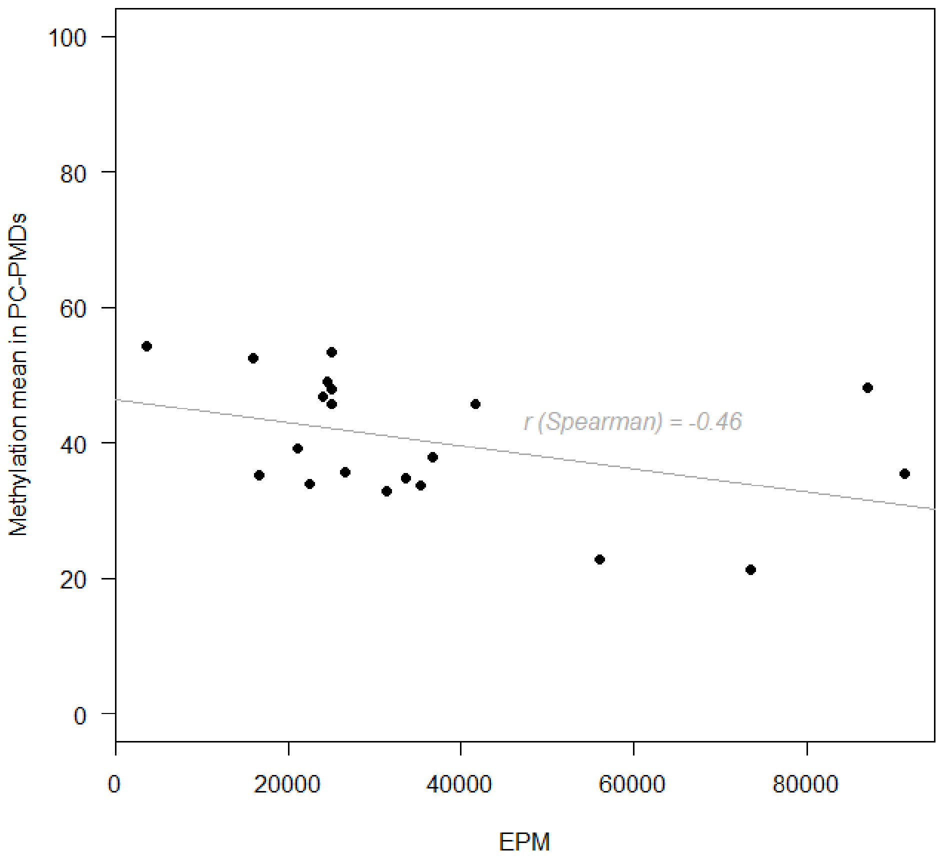


**Figure S23 |** Correlation between EPM and DNA methylation in PC-PMDs.

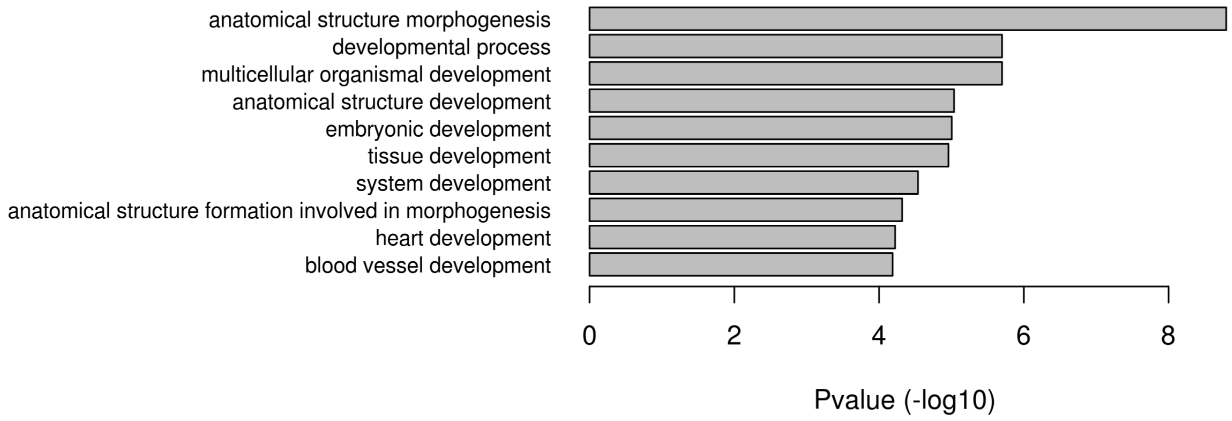


**Figure S24 |** Gene set enrichment analysis of genes from the list in the supplementary Table 7 [42].

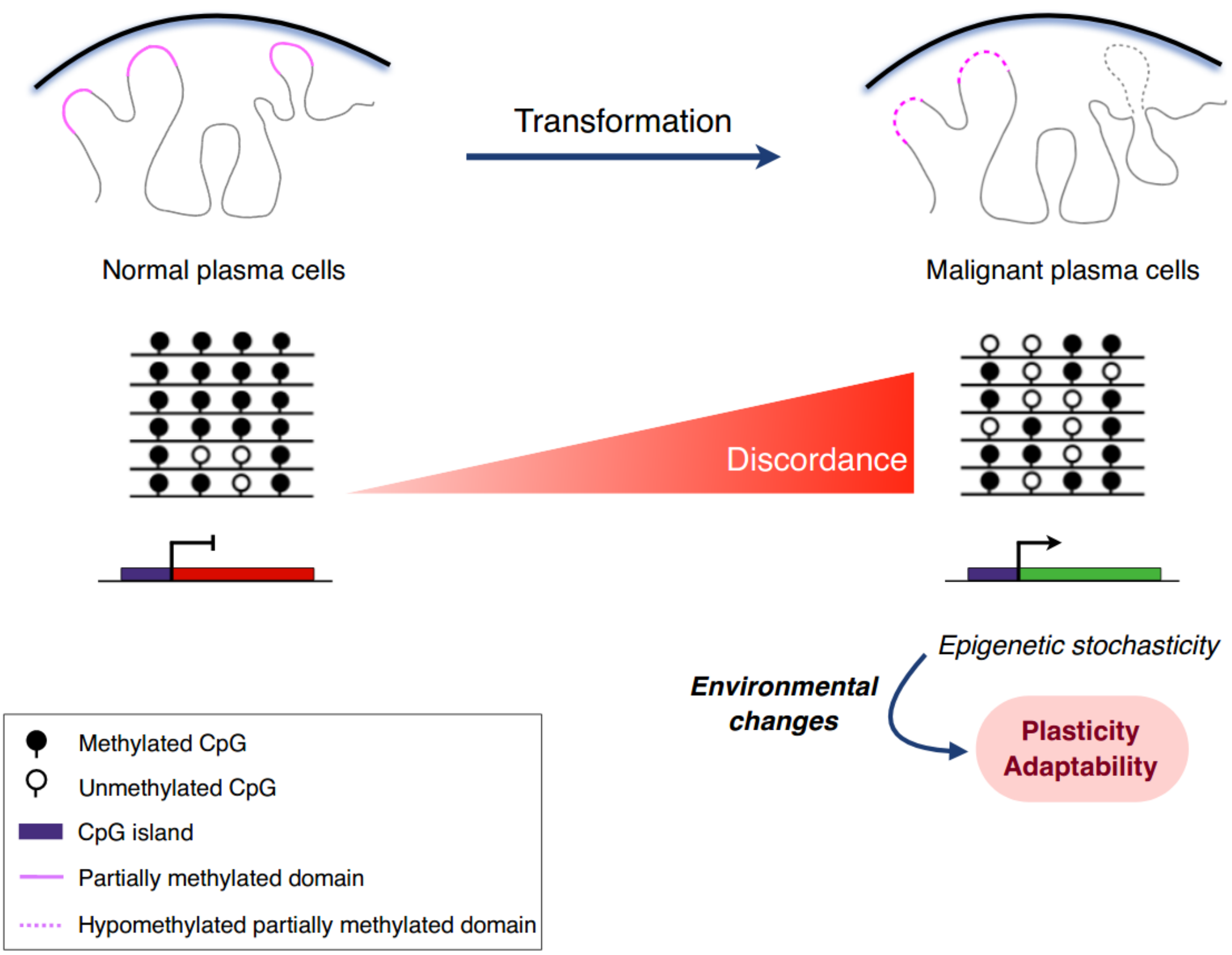


**Figure S25 |** A proposed model of the role of PMD instability in MM onset and development. Malignant transformation is accompanied by increased hypomethylation of PMDs which contributes to epigenetic, transcriptomic and 3D organization heterogeneity across patients. The lack of accurate global DNA methylation maintenance also drives intrapatient DNA methylation heterogeneity which contributes to a decoupling relationship between promoter methylation and gene expression (genes with methylated promoter but not repressed) and transcriptomic variability. We therefore hypothesize that this epigenetic stochasticity could provide a selective advantage to the tumor cell by increasing its plasticity and adaptation to environmental changes.
